# Statistics of spatial rotations in 3D electron cryo-microscopy by unit quaternion description

**DOI:** 10.1101/733881

**Authors:** Mingxu Hu, Qi Zhang, Jing Yang, Xueming Li

**Author notes:** These authors contributed equally to this work (Mingxu Hu, Qi Zhang). Correspondence should be addressed to X.L. and J.Y.

## Abstract

Electron cryo-microscopy (cryoEM) involves the estimation of orientations of projection images or three-dimensional (3D) volumes. However, the lack of statistical tools of rotations in cryoEM fails to answer the growing demands for adopting advanced statistical methods. In this study, we develop a comprehensive statistical tool specialized for cryoEM based on an unit quaternion description of spatial rotations. Some basic properties and definitions of the quaternion, as well as a way to use the unit quaternion to describe and perform rotations, are first recalled. Then, based on the unit quaternion, the distance and geodesic between rotations are designed for cryoEM to enable comparisons and interpolations between rotations, which are prerequisites of statistics of rotations in 3D cryoEM. Further, methods of directional statistics specialized for cryoEM are developed, including calculations of the average rotation, sampling, and inference with uniform and angular central Gaussian (ACG) distribution, as well as an estimation of the rotation precision. Finally, the method of handling molecular symmetry is introduced. Using the unit quaternion system for cryoEM, we provide comprehensive mathematical tools for the analysis of spatial rotations in cryoEM.

## 1 Introduction

In the three-dimensional (3D) reconstruction of electron cryo-microscopy (cryoEM), particle images or pre-reconstructed sub-tomograms of biological samples need to be rotated in 3D spaces to specific orientations in order to enable the reconstruction of a 3D density map[1, 2, 3, 4, 5, 6, 7]. Accordingly, the description, determination, operation and analysis of rotations are important for cryoEM. Different algorithms are being developed to determine the rotation, i.e., orientation, parameters in order to achieve a better rotation precision for higher resolution. There are growing demands for adopting advanced statistical methods in the development of such algorithms. In this study, we develop comprehensive statistical tools specialized for cryoEM to meet these demands.

A proper system of spatial rotation description is required for such statistical tools. To describe spatial rotations, three approaches have been widely used in different fields, namely the rotation matrix, Euler angles, and unit quaternion. The rotation matrix is the most basic one, and can be directly used to rotate a 3D vector or object. However, nine parameters with non-linear constraints in a 3 × 3 matrix are required to describe a 3D rotation, which is often infeasible for algorithms to directly estimate and optimize them. The Euler angle description is an alternative approach that parameterizes the rotation matrix using three angle values that represent sequential rotations around three given axes, for example, the *Z, Y*, and *Z* axes. Euler angles are intuitive, and they have therefore been widely adopted in cryoEM for more than 40 years[1, 2, 3, 4, 5, 6, 7]. However, there are several drawbacks with the Euler angle description, such as the non-uniqueness of the rotation description, dependency on the choice of the rotation axes, the gimbal lock issue, and the scarcity of sampling and statistics approaches in the spatial rotational space.

The unit quaternion is a different system from the rotation matrix and Euler angles, and overcomes most drawbacks of Euler angles as mentioned above [8]. Several packages in cryoEM, including SPIDER[7] and EMAN2[5], have began to use the unit quaternion to perform basic rotations. There are more advantages in using the unit quaternion comparing with Euler angles, which are still not well explored in cryoEM, for example, the statistical analysis of rotations, including averaging, interpolation, sampling, and inference. The use of such features should not only enhance the data analysis in the traditional cryoEM method, but also enable the applications of the modern statistical inference method, such as the particle filter algorithm, which has been implemented in THUNDER[9].

The present work aims to develop specialized statistical tools for cryoEM to enable statistical analysis of rotations based on unit quaternion description. As the majority of the readers may be unfamiliar with the quaternion, Section 2 gives a brief recall of definitions and basic properties of the quaternion, as well as a way to use the unit quaternion to describe and perform rotations. In Section 3, the distance and geodesic between two rotations are defined, which are specialized for cryoEM. These serves as the prerequisites of the statistics tool development. Further, in Section 4, the statistical methods for performing sampling and numerical analysis in the rotational space of cryoEM are introduced and developed. Finally, in Section 5, the method of handling molecular symmetry and corresponding space division of asymmetric units are introduced.

## 2 Background of the quaternion and unit quaternion description

The quaternion and unit quaternion description of rotation has been developed and applied in different fields. In these section, we recall some of the properties and features of the quaternion and unit quaternion, which is potentially suitable for cryoEM. For more general introductions, we refer to these references[10, 11].

Notations in this article are specified according to the following rules. The 3D vector is denoted using a letter with a bold font and an arrow on hat, as in 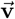. The corresponding unit vector is decorated with a triangle hat, as is the case with 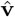. The quaternion in vector form and a four-dimensional (4D) vector are denoted using a letter with a bold font, as in **q**. The spatial rotation operator is denoted as *R*. All vectors are presented as column matrices, and the superscript, *T*, indicates matrix transpose. Moreover, the 3D space is denoted as 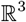, and all unit 3D vectors form a 2D surface of a unit sphere denoted as *S*^2^ in 3D space. Similarly, the 4D space is denoted as 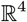, and all unit 4D vectors form a 3D hypersurface of the unit sphere denoted as *S*^3^ in 4D space. The set containing all 3D spatial rotations is denoted as *SO*(3).

### 2.1 Definitions and basic properties of the quaternion

A quaternion is an extension of a complex number. A quaternion **q** is defined as a hypercomplex number composed of a real part and three imaginary parts as

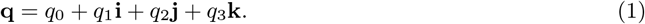

Equivalently, the quaternion is often written in 4D vector format as

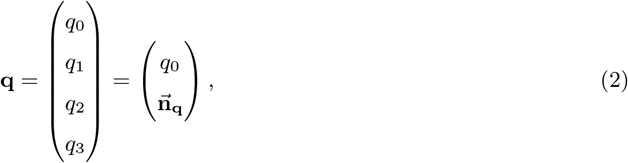

where **{i, j, k}** are three imaginary units, and 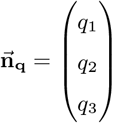.

The conjugate of a quaternion is defined as

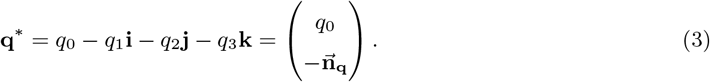

The norm of a quaternion **q** is defined as

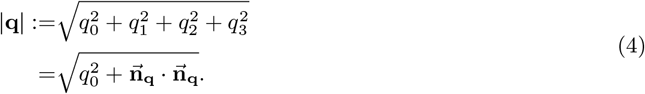

The multiplication of two quaternions, **q** and **q′**, is calculated based on the multiplication rule of their basis elements, 1, **i, j**, and **k** as

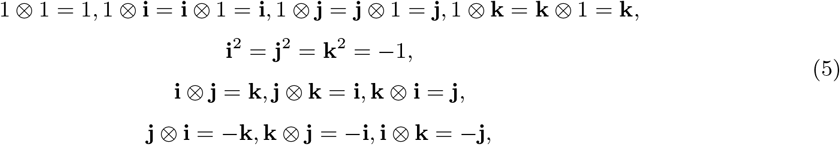

where ⊗ represents the quaternion product. The result of quaternion multiplication can also be written in vector format as

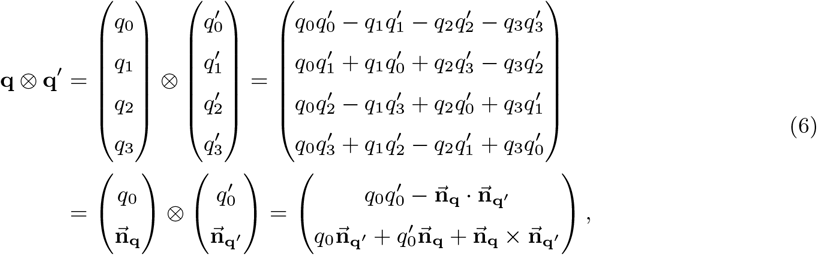

where · and × represent the inner product and cross product of vectors. The complex form and vector form of quaternion multiplication are equivalent. For simplicity, the vector form of a quaternion is frequently used as follows. More basic properties of quaternion algebra are listed in Appendix I.

### 2.2 Unit quaternion description of three-dimensional spatial rotation

If norm equals to 1, i.e., |**q**| = 1, the quaternion **q** is called a unit quaternion. The product of a unit quaternion **q** with its conjugate is 1, as **q** ⊗ **q\*** = **q\*** ⊗ **q** = 1; all unit quaternions form *S*^3^ (see Appendix I for more properties). The unit quaternion can be used to perform 3D rotation. A brief introduction is as follows.

The rotation of a 3D vector 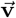 to another vector 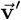 is performed using a 3 × 3 orthogonal matrix **M** with a determinant of 1, as follows.

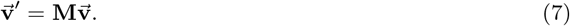

The same operation can also be achieved by performing unit quaternion multiplication. Viewing 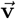 as 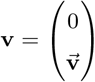 in quaternion form, the rotation operation induced by unit quaternion **q** is defined as[10]

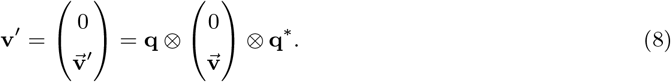

In the following context, we will not distinguish between **v** and 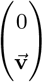. For simplicity, *R*_**q**_ is used to represent the rotation by unit quaternion **q** in the present work, i.e.,

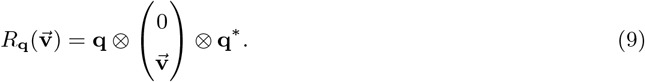

Note that a pair of unit quaternions is involved in the multiplication; hence, **q** and **−q** will not cause a different rotation[10]. Consequently, *R*_**q**_= *R*_−**q**_.

The unit quaternion explicitly represents a 3D spatial rotation around an axis. Any unit quaternion **q** can be written in the form[11]

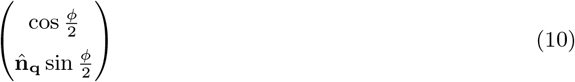

which presents a rotation around axis 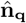 with rotation angle *ϕ*.

For sequential rotations, such as *R*_**q**_1__ followed by *R*_**q**_2__, the unit quaternion of the overall rotation is calculated as[10]

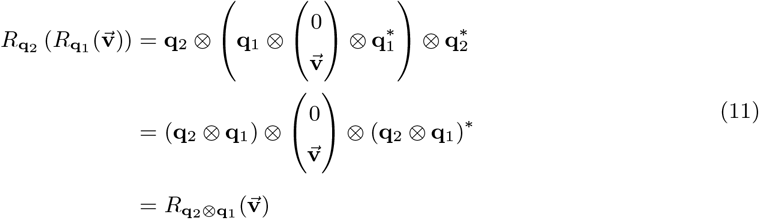

demonstrating that **q**_2_ ⊗ **q**_1_ represents the overall rotation. Moreover, the dot product and cross product of vectors are preserved under spatial rotations, i.e., quaternion multiplications (see Appendix II). This is useful for combined computations of quaternion multiplication and vector multiplication.

Finally, the conversions between three rotation descriptions, namely the rotation matrix, Euler angles, and unit quaternion description, are well-established. The formulas of these conversions are summarized and introduced in Appendix III and Appendix IV.

### 2.3 Swing-twist decomposition

If using a directional vector to represent the orientation of a 3D object, the 3D rotation of this object can be decomposed into the product of two rotations, a swing which changes the directional vector to another direction and a twist around the directional vector, which is called swing-twist decomposition[12]. This is potentially useful for rotation analysis and the application of constraints to rotation.

To demonstrate the potential application of the swing-twist decomposition in cryoEM, we consider a 2D experimental image in single-particle cryoEM, which will be rotated using unit quaternion **q** to match the projection of a 3D object. In cryoEM, we usually assume that the experimental image initially lies in the *xy* plane with norm vector 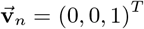. The rotation can be performed in two steps.

First, the image is rotated so that its norm vector 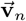 is changed to be along the projection direction 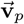, which is called the swing. 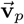 is calculated as follows.

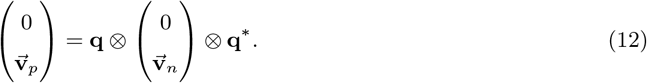

The swing rotating 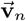 to 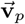 can be obtained by a rotation around the axis

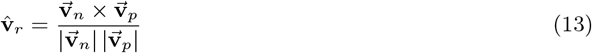

for angle

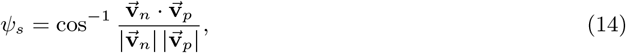

which is called the swing angle. Accordingly, the swing quaternion is

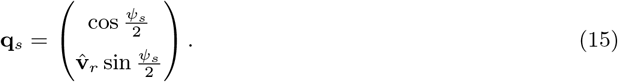

Second, the image is in-plane rotated around its norm vector; currently, 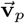, for angle *ψ_t_* to match he projection of the object, called the twist. Then, the twist quaternion is

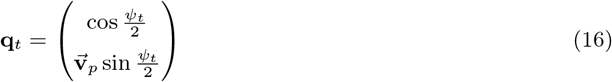

as 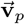 is a unit vector.

Based on the definition of swing-twist decomposition, i.e., **q** = **q**_*t*_ ⊗ **q**_*s*_, the twist quaternion 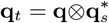 can be calculated. By comparing **q**_*t*_ with Equation (16), the twist angle *ψ_t_* can be calculated.

The decomposing **q** = **q**_*t*_ ⊗ **q**_*s*_ mentioned above is known as the swing-before-twist form. The swing-twist decomposition also has a twist-before-swing form, where the twist is performed before the swing. Because 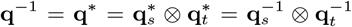, the twist-before-swing form can also be considered as the inversion of the swing-before-twist form.

## 3 Distance and geodesic between rotations in cryoEM

In cryoEM, it is important to compare and evaluate the differences between rotation parameters. However, a quantitative measure for such difference is still missing in cryoEM. Here, we used the concept of the distance between two rotations based on the unit-quaternion system as a quantitative measure.

More importantly, the definition of the distance is a prerequisite of directional statistics in 3D rotational space *SO*(3), which enables statistical analysis, i.e., sampling and inference, in the rotation parameter space (discussed in the following sections). In some cryoEM programs, such as RELION[1] and cryoSPARCs[2], HEALPix[13] grid distance under constrained conditions with the Euler angle system has been used as an approximate measure for the differences of rotation parameters. However, the properties of Euler angles limit the application of the rotation distance. In contrast, the definition of the distance and corresponding geodesic based on the unit quaternion has shown its advantages as a general mathematical tool for angular analysis and rotational operations.

### 3.1 Definition of distance between rotations

The angle between two 3D unit vectors 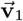 and 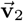 can be defined as the distance between them as follows.

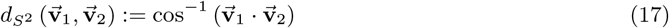

which ranges [0,*π*]. Because 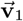 and 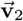 correspond to two points on the unit sphere *S*^2^, the distance between 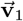 and 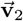 is the length of the shortest arc between their corresponding points on *S*^2^. The use of the angle value as the distance is straightforward for orientation analysis. However, the unit quaternion presents a rotation rather than a 3D vector. Thus, we cannot directly define an angle value to quantify the distance between two rotations.

There are several existing definitions related to the distance between rotations in *SO*(3), as summarized by Du et al.[14]. However, these definitions do not fit well into a cryoEM scenario as none of these definitions are related to attributes of interest in cryoEM. Thus, we raise the definition of distance between two rotations in *SO*(3), *R*_**q**_1__ and *R*_**q**_2__, as

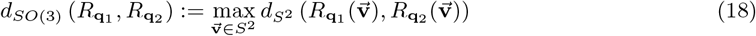

which is the maximum distance between two 3D vectors, 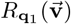 and 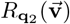, generated by rotating an arbitrary unit vector 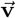 using unit quaternion **q**_1_ and **q**_2_. By abuse of notations, we denote *d*_*SO*(3)_ (*R*_**q**_1__, *R*_**q**_2__) as *d*_*SO*(3)_ (**q**_1_, **q**_2_) in the following context. This definition meets all the requirements of the distance definition in mathematics (see Appendix V-I).

This distance can be calculated as

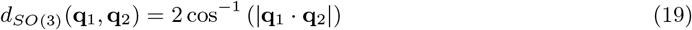

(see Appendix V-II), where the coefficient 2 in the beginning of right item of Equation (19) is a byproduct of the fact that unit quaternion space *S*^3^ and rotation space *SO*(3) form a 2 – 1 homomorphism. The absolute value in this calculation indicates that **q**_*i*_ and −**q**_*i*_ (*i* = 1, 2) represent the same rotation and will not cause different distances.

In the cryoEM scenario, the distance we defined above provides a simple estimate of the possible maximal changes of projection directions resulting from different rotations as

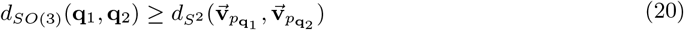

where 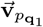 and 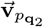 are the projection directions that result from rotations *R*_**q**_1__ and *R*_**q**_2__, respectively (see Appendix V-III). For example, a particle image in single-particle cryoEM may be rotated for multiple rounds to search for its final projection direction. The distance between rotations in adjacent rounds gives direct information about whether the search has become stable.

### 3.2 Geodesic between rotations

In geometry, the geodesic is a curve or line indicating the shortest path between two points in a surface. Here, we introduced the geodesic between two rotations, *R*_**q**_1__ and *R*_**q**_2__, i.e., as the evolvement from the rotation *R*_**q**_1__ to the rotation *R*_**q**_2__ along the shortest path in *SO*(3), which was then used for the interpolation between two rotations.

In the unit quaternion system, this geodesic is the shortest arc connecting the corresponding unit quaternions of rotations *R*_**q**_1__ and *R*_**q**_2__ on the surface *S*^3^. Note that the distances between **q**_1_ and **q**_2_, and between **q**_1_ and −**q**_2_ are the same. However, the shortest arcs connecting **q**_1_ and **q**_2_, and connecting **q**_1_ and −**q**_2_ may have different lengths. It can be easily determined that the shortest of the two arcs has a length equal to half of the distance of corresponding rotations. Thus, for clarity and to match the definition of the distance mentioned above, only the shortest arc was selected as the geodesic. Srivastava et al.[15] defined and discussed the same geodesic. However, their definition of the rotation distance is different from ours based on Equation (18), i.e., half of the length of our distance definition (Equation (19)).

Based on the definition of the geodesic, Srivastava et al. developed a method of spherical linear interpolation (SLERP) for 3D rotations using a unit quaternion. Below, we briefly review their conclusion, which is considered as an example of the geodesic application.

The geodesic between *R*_**q**_1__ and *R*_**q**_2__ can be described as a function

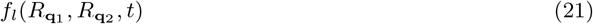

of an affine parameter *t* ∈ [0,1]. By abuse of notations, in the following context, we denote *f_l_*(*R*_**q**_1__, *R*_**q**_2__, *t*) as *f_l_*(**q**_1_, **q**_2_, *t*), and the value of this function is also a unit quaternion representing the rotation. This function describes the relation distance from *R*_**q**_1__, i.e.,

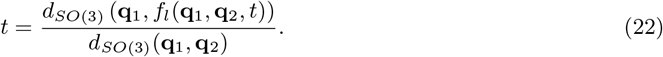

Therefore, it is easy to obtain

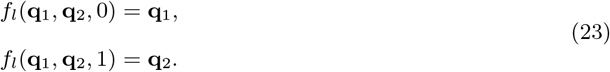

Based on the geometry of the geodesic, the analytical formula of the geodesic function can be derived as

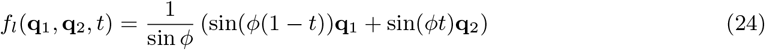

when **q**_1_ · **q**_2_ ≥ 0, and

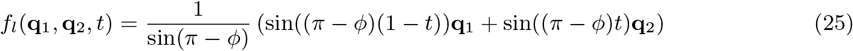

when **q**_1_ · **q**_2_ < 0, where *ϕ* = cos^−1^ (**q**_1_ · **q**_2_), ranges [0,*π*].

The formula can be used to calculate interpolations between two rotations. As an example, we calculated the interpolation between two rotations *R*_**q**_1__ = (−0.884, −0.265, −0.123, 0.364)^*T*^ and *R*_**q**_2__ = (0.370, 0.424, −0.686, −0.461)^*T*^ (Figure 1a), and converted the interpolated rotations to Euler angles with a ZYZ convention used in cryoEM (Figure 1b). The plots show that the changes of three Euler angles *ϕ, θ*, and *ψ*, are not linear, compared with the linear property of the distance from *R*_**q**_1__ to the interpolated rotation. Conversely, this implies that the linear interpolation of Euler angles will not yield “linear” changes of the rotations. In fact, it is very difficult to define a geodesic in the Euler angle system.

**Figure 1:**
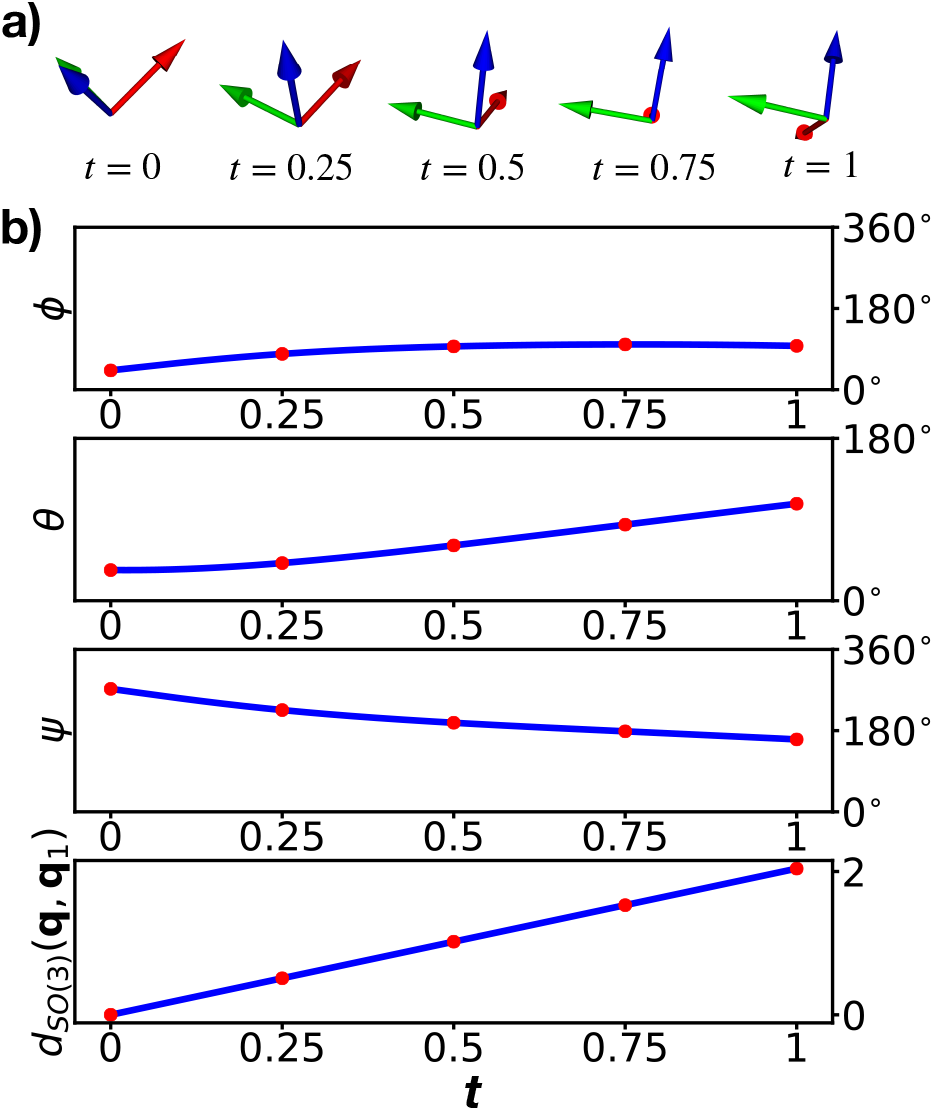
Geodesic between two rotations. Geodesic between two rotations, **q**_1_ = (−0.884, −0.265, −0.123, 0.364)^T^ and q_2_ = (0.370, 0.424, −0.686, −0.461)^T^. **a**, Snapshots of *x*-axis, *y*-axis, and *z*-axis rotated by rotations along the geodesic, where the red arrow indicates the *x*-axis, the green arrow indicates the *y*-axis, and the blue arrow indicates the *z*-axis. **b**, ZYZ Euler angle description and the distance to **q**_1_ of rotations along the geodesic. Red dots represent the snapshots in **a**.

## 4 Statistics of rotations in cryoEM

Sampling and randomization in the rotational space is crucial for 3D reconstruction in order to avoid the so-called local optimization problem or to statistically cover a large rotational space using a limited number of samples. In the Euler angle system, there has been widespread use of grid sampling by setting a step size for each of three Euler angles. However, the rotational space is not a linear space, and equal-step sampling of the Euler angle will not generate uniform coverage in rotational space; hence, it is easy to cause bias. The HEALPix[13] method is an improved algorithm for approximately uniform sampling on a rigid grid with a number of grid points proportional to *N*^3^, where *N* is a positive integer and indicates the precision of the grid interval. The drawback of these Euler-angle-based methods is that it is difficult to generate samples with a real uniform distribution or more complicated distribution. The increased number of applications of the statistical inference algorithm in cryoEM, such as the particle filter based on random importance sampling, has increased the demand for better mathematical tools to perform random sampling in rotational space.

In general, there are two types of requests for random sampling. One is to generate samples in rotational space with a specific statistical distribution. Another one is to estimate the statistical properties for a series of given rotation parameters. Corresponding methods and theories are usually referred to as directional statistics, and the unit quaternion has shown advantages for such purposes. Together with the distance and geodesic defined above, we introduce and developed tools of unit quaternion for cryoEM. Here, we first introduce a method to calculate the average of a set of rotations. Second, we discuss the method of uniform sampling in SO(3). Third, we introduce the angular central Gaussian (ACG) distribution. Finally, a specific ACG distribution is shown as an example to demonstrate a costume application of ACG distribution. Based on such distribution, we derived a calculable formula that should also be useful in the estimation of the angular precision and angular stability of a particle during 3D alignment.

### 4.1 Average rotations

The average of a series of rotations is one of the simplest statistical parameters, and it is useful when generating ACG distribution (discussed later). Based on the definition of the distance in Equation (18), we defined the geometric mean[16] as the average of a given set of rotations {*R*_**q**_*i*__}_1≤*i*≤*N*_ with corresponding weights {*w_i_*}_1≤*i*≤*N*_ for each rotation,

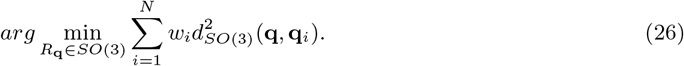

However, it is difficult to calculate Equation (26)[16]. To avoid this problem, we proved that the geometric mean in Equation (26) can be approximated using the projective arithmetic mean[17] (see Appendix VI). The projective arithmetic mean of rotations using unit quaternion was introduced both by Horn[18] and by Markley et al.[19], which was the normalized principal eigenvector of the matrix

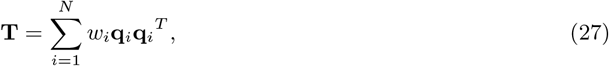

where 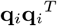 is a tensor of **q**_*i*_ and itself. Eigenvalues and eigenvectors can be determined using the linear algebra method. While only the principal eigenvector is needed, we usually use the iterative formula[20]

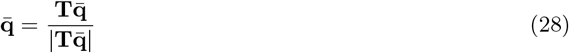

to approximate the principal eigenvector. Equation (28) will not converge only when the largest two eigenvalues of **T** are strictly equal, which nearly never happens in numerical calculations.

### 4.2 Uniform distribution in *SO*(3)

Uniform sampling in rotational space is frequently used in cryoEM alignment to search for the global optimal solution of molecular orientations. The unit quaternion offers a simple way to generate uniform distributed unit quaternions {**q**_*i*_}_1≤*i*≤*N*_ on *S*^3^, representing uniform distributed rotations {*R*_**q**_*i*__}_1≤*i*≤*N*_ in *SO*(3)[21].

Using a 4D Gaussian distribution *N*_4_(0,**I**) with a mean of zero and covariance matrix of a 4 × 4 identity matrix **I**, a series of 4D vectors can be sampled. Then, by normalizing these vectors with norm 1, we obtain unit quaternions represented by **q** uniformly distributed in *S*^3^, i.e.,

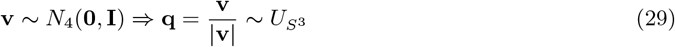

where **v** is a 4D vector sampled from *N*_4_(0, **I**), and *U*_*S*^3^_ denotes the uniform distribution in *S*^3^. Accordingly, the corresponding rotation *R*_**q**_ obeys a uniform distribution in *SO*(3). As an example to mimic the global search in cryoEM, we generated a set of rotations with uniform distribution, and applied them on an initial orientation along the *Z*-axis. Both the resulting projection directions and in-plane rotational angles by swing-twist decomposition follow a uniform distribution (Figure 2a and Figure 2b).

**Figure 2:**
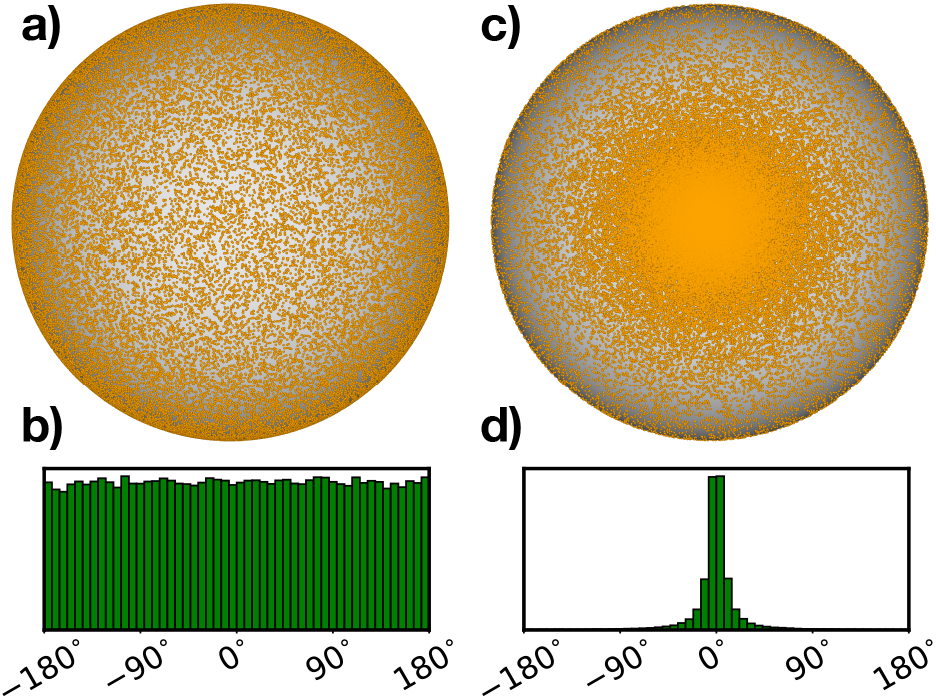
Uniform distribution and ACG distribution. 100,000 rotations are sampled from the uniform distribution and ACG distribution with covariance matrix **A** = *diag*(20^2^, 1,1,1), respectively, and are applied to an initial direction (0, 0,1)^*T*^. **a** and **b** are respectively plots of the projection directions and in-plane rotation angles from rotations with a uniform distribution. **c** and **d** are respectively plots of the projection directions and in-plane rotation angles from rotations with an ACG distribution. The swing-twist decomposition was used to derive the projection directions and in-plane rotation angles.

### 4.3 Statistics obtained using ACG distribution

In the local search of cryoEM 3D alignment, we often expect to perform intensive searches around a given orientation/rotation with high probability. As the distance from the given rotation increases, the probability of finding the correct solution is decreased, and hence fewer searches are needed. Under such a situation, sampling with a Gaussian distribution is often performed. However, the rotational space is not a linear space, and the Gaussian distribution with a requirement of a linear space is not satisfied. Alternatively, the ACG distribution[22] provides a solution in rotational space with features similar to that of a Gaussian distribution. In general, the ACG distribution is a useful tool in rotational space to generate or analyze samples with a maximal central distribution. THUNDER[9] has used tools of ACG distribution to perform sampling in rotational space.

#### 4.3.1 Definition of ACG distribution

Similar to the uniform distribution, the generation and normalization of 4D vectors from Gaussian distribution *N*_4_(**0, A**) with zero mean but with a no-identity covariance matrix **A** that is positive definite and symmetric leads to unit quaternions that are represented by **q**, obeying the ACG distribution as[22]

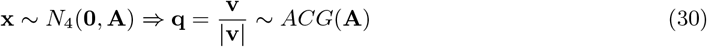

where **v** is a 4D vector sampled from *N*_4_(**0, A**). The probability density function of the ACG distribution is[22]

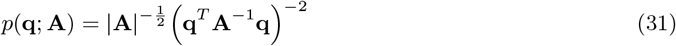

which reaches a maximum when **q** equals the principal eigenvector of the covariance matrix **A**[22]. Therefore, by selecting a covariance matrix **A** with a principal eigenvector along **q**, we can obtain an ACG distribution with maximum probability density at **q**. As an example, Figure 2c and Figure 2d demonstrate the distribution of the projection direction and in-plane rotational angles derived from the swing-twist decomposition, respectively, where **A** is a diagonal matrix with principal eigenvector along (1, 0, 0, 0)^*T*^. A general method is introduced to generate such a covariance matrix (see Appendix VII).

Given a set of unit quaternions {**q**_*i*_}_1≤*i*≤*N*_, the estimation for the covariance matrix **A** is useful for the probability analysis or generating new random rotations. While missing an explicit maximum-likelihood method to estimate **A**, an iterative formula[22] is usually used to approximate **A**

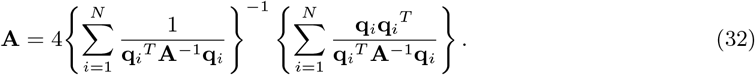

#### 4.3.2 Local perturbation and confidence area

As mentioned before, a series of rotations {*R*_**q**_*i*__}_1≤*i*≤*N*_ can be generated using an ACG distribution around a given rotation 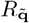 by selecting a covariance matrix **A**. Here, we used a special form of covariance matrix to generate rotations, which can be conveniently correlated with the angular accuracy of the local search.

To obtain {*R*_**q**_*i*__}_1≤*i*≤*N*_, we first generate a set of unit quaternions 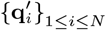 with an ACG distribution *ACG*(*A*), where a special covariance matrix

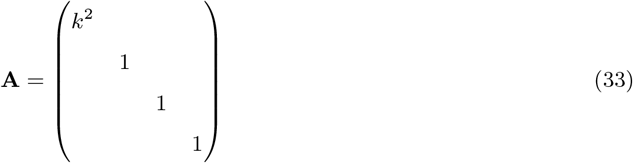

is chosen with *k* > 1 so that the principal eigenvector of A is along **q_e_**= (1, 0, 0, 0)^*T*^, *R*_**q_e_**_ is a still rotation, i.e., no rotation. Accordingly, 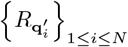 present perturbations for the still rotation. By appending these generated rotations to a given rotation 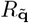, i.e., 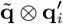 we obtain perturbations {*R*_**q**_*i*__}_1≤*i*≤*N*_ for 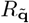 as

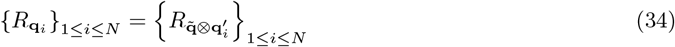

which still obeys a specific ACG distribution.

In the above covariance matrix **A**, the parameter *k* is used to regulate the range of perturbations. A larger *k* value leads to a narrower distribution of {**q**_*i*_}_1≤*i*≤*N*_ in *S*^3^, i.e., perturbations over a smaller range. Clearly, for the local search during cryoEM 3D alignment, the range of perturbations is directly correlated to the angular accuracy of the search. In the following, we develop the relationship between *k* and the angular accuracy of the local search, which may be used to generate a perturbation with a given accuracy requirement.

Considering an arbitrary unit vector 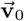 and a resulting unit vector 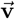 from rotating 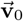 by **q** ~ *ACG*(**A**) with **A** in Equation (33), the inner product of 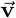 and 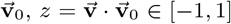, equals the cosine of their angular distance. The angular distance, i.e., cos^−1^(*z*), reflects the intensity of the perturbation obtained by **q**.

To characterize the intensity of the perturbation, we first derived the probability density of the resulting unit vector 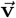, which is (see Appendix VIII)

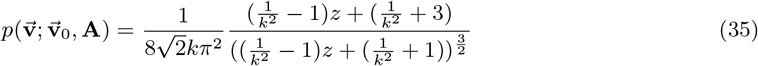

where 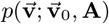 reaches a maximum when 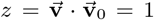, i.e., 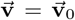, and 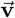 obeys a distribution invariant under rotation around 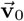 (Appendix VIII, Figure 3a). Then, we introduced the concept of the confidence area [*z*_0_, 1] with confidence level *a*, which satisfies

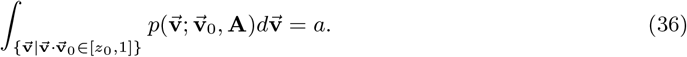

**Figure 3:**
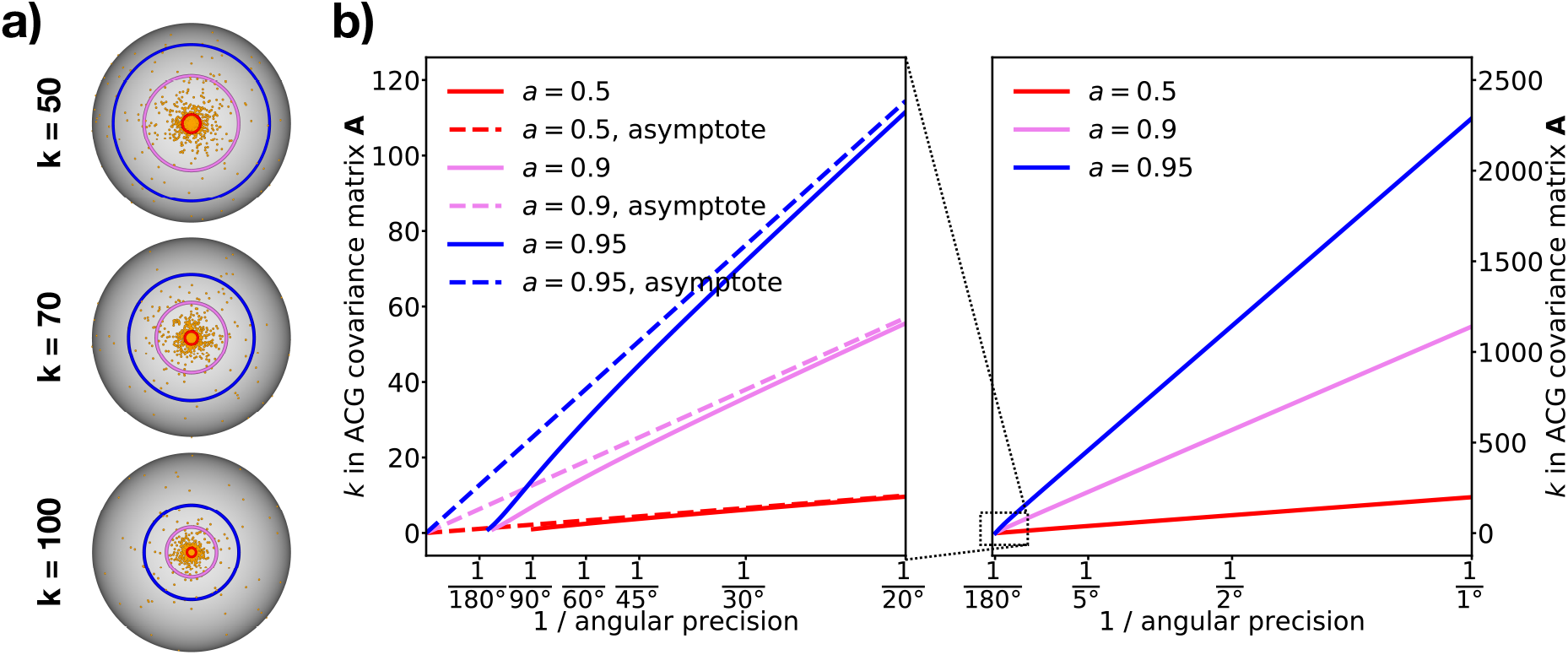
ACG distribution with different covariance matrix parameter *k*. **a**, distributions with *k* equals to 50, 70, and 100. In each plot, 3000 rotations are sampled from an ACG distribution with a covariance matrix of *diag*(*k*^2^,1,1,1). The resulting vectors rotated from (0, 0,1)^*T*^ are plotted as orange points. Boundaries of the confidence area with multiple confidence level *a* of 0.5, 0.9, and 0.95 are shown as red, violet, and blue circles, respectively. **b**, The correlation between *k* in the ACG covariance matrix **A** and the reciprocal of the angular precision. Multiple confidence levels are plotted. Dashed lines indicate the asymptotes of this correlation.

The confidence level *a* describes the probability of making a perturbation up to *z*_0_, or up to the angle distance of cos^−1^(*z*_0_) (Figure 3b). In other words, the confidence level can also be considered as the percentage of samples fallen into a range with angle distance up to cos^−1^(*z*_0_). Therefore, *ϵ* = cos^−1^ (*z*_0_), can be defined as the angular precision of rotation sampling under a given confidence level, *a*. We developed the following formula to calculate *z*_0_ with given *k* and *a* (see Appendix IX)

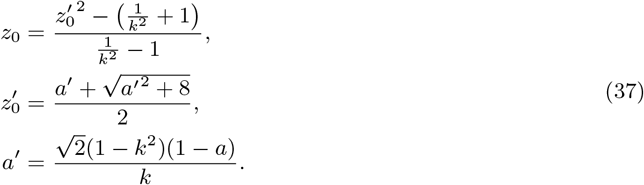

To simplify the computation, we derived an approximation formula to calculate angular precision *ϵ* from *k* and *a*, or to calculate *k* from *ϵ* and *a*, as

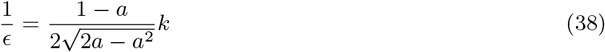

In fact, this formula is the asymptote of 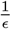 when *k* → ∞ (Figure 3b). The correctness of this approximation formula is proven in Appendix X.

#### 4.3.3 Inference based on ACG distribution

In the particle-filter algorithm implemented in THUNDER[9], the posterior probability density function of the rotation estimates is described by the distribution of a series of rotations {*R*_**q**_*i*__}_1≤*i*≤*N*_. This generates the requirement for the inference of distribution of {*R*_**q**_*i*__}_1≤*i*≤*N*_.

Considering the inverse process of the previous section, {*R*_**q**_*i*__}_1≤*i*≤*N*_ can be considered as perturbations for their average 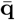. Thus, 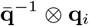 or corresponding rotations 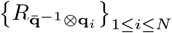 are the perturbations for still rotation **q**_*e*_ = (1, 0, 0, 0)^*T*^. For simplicity, we assume that 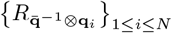, obeying the ACG distribution, and has a covariance matrix in Equation (33). Then, we developed an iterative method to determine the maximum-likelihood approximation of *k* from 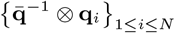, which is

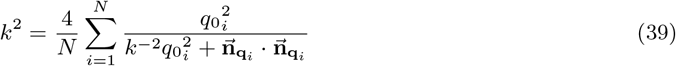

where 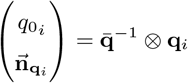. The correctness of this iterative method is proven in Appendix XI.

Combining Equation (37) with Equation (39), we developed a method to estimate the angular precision of the parameter search from a series of samples generated by the particle-filter algorithm in THUNDER[9]. As an example, we performed the ab initio 3D reconstruction of a dataset of the proteasome (EMPIAR-10025) using 112,412 particles. A global search was performed under three sequentially increasing cutoff frequencies, 56.1*Å*, 16.8A, and 9.9A. For each cutoff frequency, the average angular precision of all particles increased and converged (Figure 4, blue, red, and orange dots, respectively). During a local search, as the cutoff frequency increased in each round from 9.9*Å* to 2.69*Å*, the average angular precision rapidly increased from 1.42° and then converged at 0.704° (Figure 4, violet dots). Finally, defocus refinement was performed. As the resolution increased from 2.47*Å* to 2.33*Å*, the angular precision was improved to 0.661° (Figure 4, blue dots).

**Figure 4:**
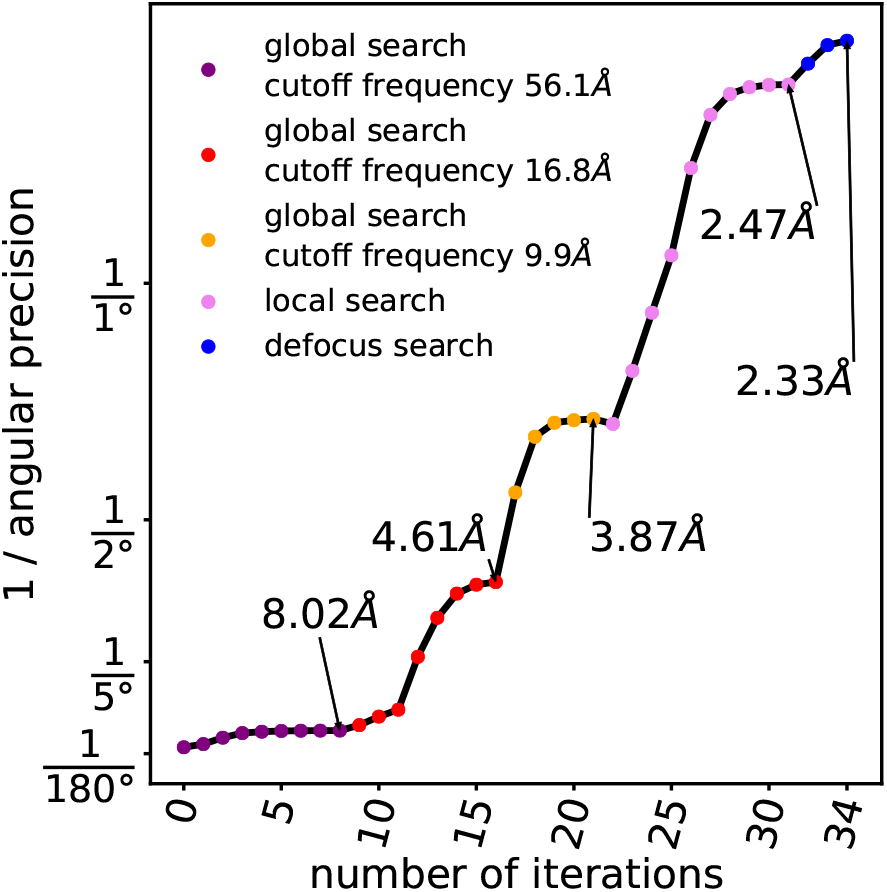
The angular precision increases as the refinement is iterated. 3D refinement is performed using a proteasome dataset (EMPIAR-10025) with 112,412 particles and an ellipsoid as initial model. The reciprocal of the average angular precision is plotted against the iteration number of refinement. A global search is performed under three sequentially increasing cutoff frequencies, followed by a local search and a defocus search. The resolutions of the refining reference at several iterative rounds are labeled.

## 5 Molecular symmetry by unit quaternion

In cryoEM 3D reconstruction and alignment, molecular symmetry is an important property that can be used to enhance the signal and reduce the computation. In this section, we show how to apply unit quaternions to simplify calculations of molecular symmetry.

As molecules are chiral in cryoEM, there are only five classes of molecular symmetry groups. These are the cyclic group (*C_n_*), dihedral group (*D_n_*), tetrahedral group (*T*), octahedral group (*O*), and icosahedral group (*I*). With unit quaternions, the symmetric operations of these symmetry groups can be calculated conveniently, from no more than two generators, which were introduced by Conway et al.[11]. Based on Conway’s method, we derived the generators of these molecular symmetry groups under the orientation conventions in cryoEM (Table 1). The cyclic group is the simplest one, and has only one generator. The other four groups need two generators.

**Table 1:** Molecular symmetry groups.

| Symmetry Group | Symbol | Generators | Axis of Generators | Number of Elements |
| --- | --- | --- | --- | --- |
| Cyclic | $C_n$ | $\left(\cos \frac{\pi}{n}, 0, 0, \sin \frac{\pi}{n}\right)^T$ | $C_n$ on $Z$ | $n$ |
| Dihedral | $D_n$ | $\left(\cos \frac{\pi}{n}, 0, 0, \sin \frac{\pi}{n}\right)^T$,<br>$(0, 1, 0, 0)^T$ | $C_n$ on $Z$,<br>$C_2$ on $X$ | $2n$ |
| Tetrahedral | $T$ | $\left(\frac{1}{2}, 0, 0, \frac{\sqrt{3}}{2}\right)^T$,<br>$\left(0, 0, \frac{\sqrt{6}}{3}, \frac{\sqrt{3}}{3}\right)^T$ | $C_3$ on $Z$,<br>$C_2$ on $\left(0, \frac{\sqrt{6}}{3}, \frac{\sqrt{3}}{3}\right)^T$ | 12 |
| Octahedral | $O$ | $\left(\frac{\sqrt{2}}{2}, 0, 0, \frac{\sqrt{2}}{2}\right)^T$,<br>$\left(\frac{1}{2}, \frac{1}{2}, \frac{1}{2}, \frac{1}{2}\right)^T$ | $C_4$ on $Z$,<br>$C_3$ on $\left(\frac{1}{\sqrt{3}}, \frac{1}{\sqrt{3}}, \frac{1}{\sqrt{3}}\right)^T$ | 24 |
| Icosahedral | $I$ | $(0, 0, 0, 1)^T$,<br>$\left(\frac{1}{2}, 0, \frac{\sqrt{5}-1}{4}, \frac{\sqrt{5}+1}{4}\right)^T$ | $C_2$ on $Z$,<br>$C_3$ on $\left(0, \frac{\sqrt{5}-1}{2\sqrt{3}}, \frac{\sqrt{5}+1}{2\sqrt{3}}\right)^T$ | 60 |

In 3D alignment, the sampling should be performed in the entire rotational space. If molecular symmetry exists, only the sampling in an asymmetric unit is needed. However, the asymmetric unit varies under different definitions. We usually first fix a central rotation *R*_**q**_0__, and then define the asymmetric unit *D*(*R*_**q**_0__) under symmetry group *G* as

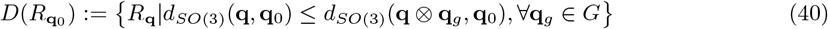

where **q**_*g*_ is a symmetric operation in *G*, and **q** ⊗ **q**_*g*_ is a symmetrically related unit quaternion of **q**. To achieve sampling in the asymmetry unit *D*(*R*_**q**_0__), we need to calculate all possible values of **q** ⊗ **q**_*g*_, and pick the one closest to **q**_0_.

An asymmetric unit is convex, which implies that the geodesic between any two rotations in an asymmetric unit lies inside the asymmetric unit. This property ensures that the SLERP interpolation between any two rotations within an asymmetric unit is still in the same asymmetric unit.

## 6 Discussion

Rotational operations and analyses are extensively used in cryoEM image analysis and 3D reconstruction. By introducing unit quaternions to replace the Euler angle system, we are able to use a more convenient and mathematically rigorous tool to operate and analysis the orientations of molecules or their projections. A disadvantage of the unit quaternion may be not as intuitional as the Euler angles in presenting the in-plane rotation and 3D orientation changes. The swing-twist decomposition provides a solution for this problem.

More importantly, the rotation operations represented by unit quaternions are tightly correlated with the 4D vector in the *S*^3^ sphere in 4D space. Accordingly, the properties of rotation can be directly derived and converted to the analysis of 4D unit vectors. This leads to the definition of the distance and geodesic. The former enables the direct comparison of rotations, and is particularly useful in the diagnosis of the orientation stability of particles during 3D alignment. The latter enables the interpolation and analysis for continuous changes of rotations, which may have potential in the analysis of flexible samples. With distance and geodesic being defined, statistics tools are established for 3D cryoEM image process. Two types of distributions, uniform and ACG, were built from the 4D Gaussian distribution of 4D vectors. These distribution models provide powerful tools of sampling and inference for either the global or local optimization of molecule orientations.

In summary, these statistics tools developed in this present work form a comprehensive system for analyzing spatial rotations in cryoEM, and they provide more freedom when processing and understanding the orientation optimization in cryoEM.

## Acknowledgements

This work was supported by funds from the National Key Research and Development Program (2016YFA0501102 and 2016YFA0501902 to X.L.), the National Natural Science Foundation of China (31722015 and 31570730 to X.L., 11471178 and 11571007 to J.Y.), Advanced Innovation Center for Structural Biology (to X.L.), Tsinghua-Peking Joint Center for Life Sciences (to X.L.), One-Thousand Talent Program by the State Council of China (to X.L.).

## Contributions

X.L. and M.H. initialized the project. X.L., M.H. and Q.Z. designed the methods. M.H., Q.Z. and J.Y. clarified mathematical issues and derived formulas. X.L. and M.H. wrote the manuscript.

## Appendix I Basic quaternion algebra property

Here, we list some basic properties of quaternion algebra. Readers may refer to[1] for more details. All of the following properties of quaternions are similar to complex numbers, except that the multiplication of quaternions is not commutative, i.e., **q**_1_ ⊗ **q**_2_ is not always equal to **q**_2_ ⊗ **q**_1_.

The inversion of a non-zero quaternion **q**, which is denoted as **q**^−1^, is defined as the quaternion that satisfies **q**^−1^ ⊗ **q** = 1 and **q** ⊗ **q**^−1^ = 1. From the rule of quaternion multiplication, i.e., Equation (6), the product of quaternion **q** and its conjugate **q\*** (Equation (3)) is

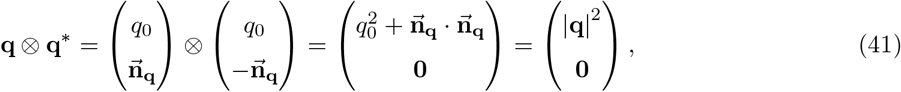

which is usually written as **q** ⊗ **q\*** = |**q**|^2^. Similarly, **q\*** ⊗ **q** = |**q**|^2^. Accordingly, **q**^−1^ can be calculated as

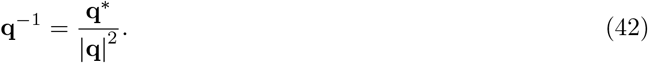

When **q** is a unit quaternion, |**q**| = 1 and **q**^−1^ = **q**^*^.

The conjugation, norm, and multiplication of quaternions satisfies the following properties.

i. (**q**\*)* = **q**.
ii. 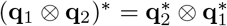.
iii. |**q**| = |**q**\*|.
iv. |**q**_1_ ⊗ **q**_2_| = |**q**_1_| · |**q**_2_|.
v. Multiplication is associative, i.e., (**q**_1_ ⊗ **q**_2_) ⊗ **q**_3_ = **q**_1_ ⊗ (**q**_2_ ⊗ **q**_3_).
vi. 1 is the multiplicative identity, i.e., 1 ⊗ **q** = **q** ⊗ 1 = **q**.

Denoting the set of all unit quaternions {**q**||**q**| = 1} as ***S***^3^, which is a 3-dimensional sphere in ℝ^4^, *S*^3^ is closed under the multiplication of quaternions, so it is a group. Moreover, *S*^3^ is a compact and Hausdorff topological subspace of ℝ^4^, and the multiplication of unit quaternions is continuous. Therefore, *S*^3^ is a compact topological group. In fact, *S*^3^ is a compact Lie group[2].

## Appendix II Dot and cross product preservation

The dot product and cross product are preserved under spatial rotation. Dot product preservation represents the property where for two 3D vectors 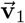 and 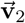, their inner product remains the same after they are rotated by the same rotation, as in

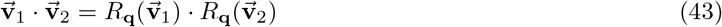

hold for all 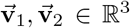 and **q** ∈ *S*^3^. Meanwhile, the cross product preservation represents the property where the rotation of the cross product of two vectors *v*_1_ and *v*_2_ equals to the cross product of the rotated vectors by the same rotation, as in

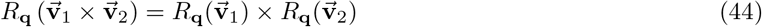

hold for all 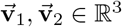 and **q** ∈ *S*^3^.

The proof is as follows. From the rule of quaternion multiplication,

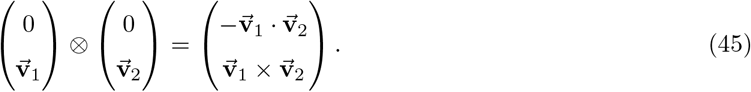

Replacing 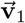 with 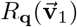 and 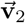 with 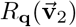 in the equation above, we obtain

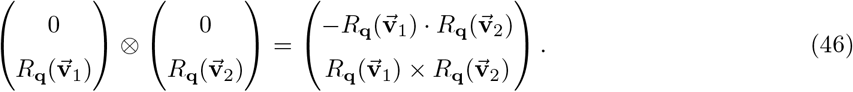

The left side of Equation (46) can be further calculated as

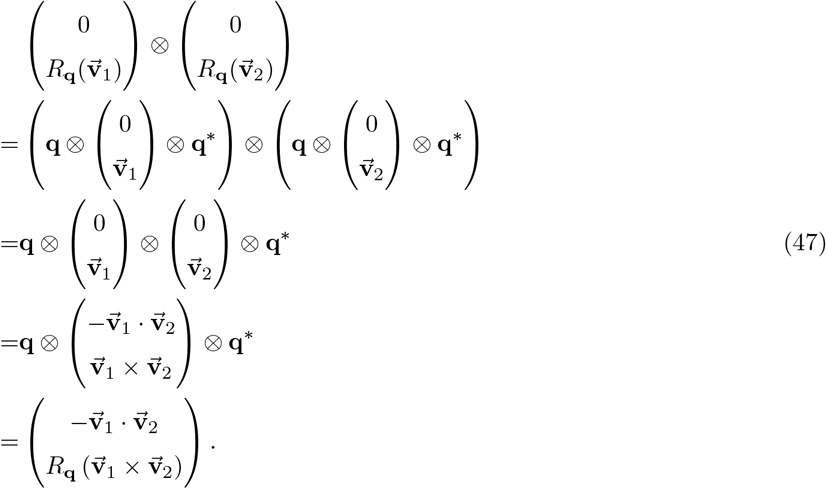

Comparing Equation (46) and Equation (47), we obtain

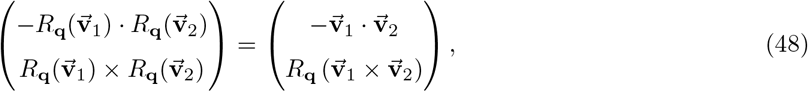

which proves Equation (43) and (44).

## Appendix III Converting unit quaternion to Euler angles

As the rotation by Euler angles *ϕ, θ*, and *ψ* in ZYZ convention is a combination of rotating *ϕ* around the *z*-axis, followed by rotating *θ* around the *y*-axis, and finally rotating *ψ* around the *z*-axis, the corresponding unit quaternion is

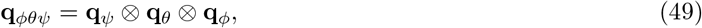

where

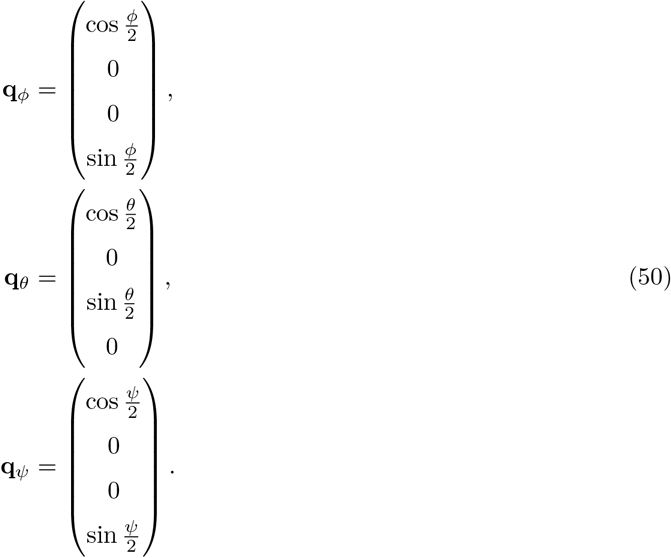

Thus, the corresponding unit quaternion **q**_*ϕθψ*_ can be calculated as

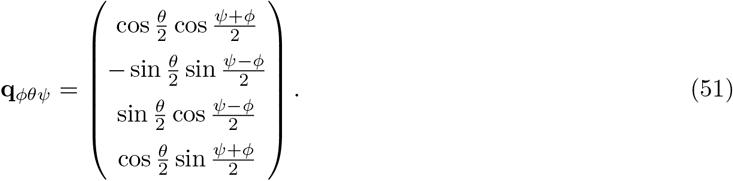

## Appendix IV Conversion between unit quaternion and rotation matrix

A3 × 3 rotation matrix **M** can be converted to the unit quaternion **q** or –**q** as follows.

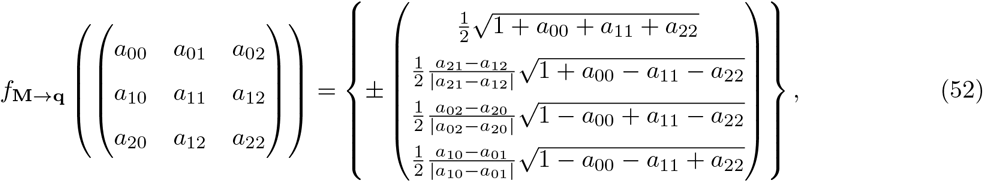

A unit quaternion **q** can be converted to a 3 × 3 rotation matrix **M** as follows.

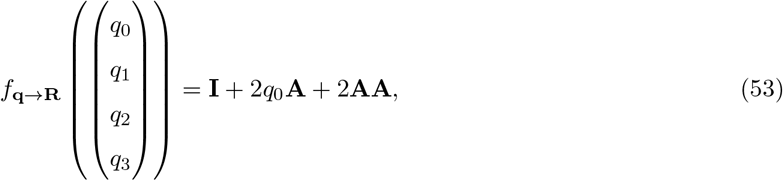

where **I** is a 3 × 3 unit matrix, and

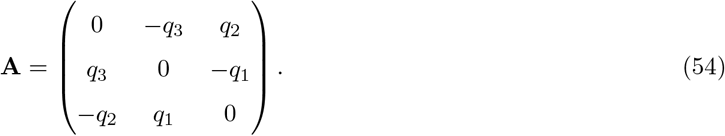

## Appendix V Distance between rotations

### Appendix V-I Proof of wellness of distance definition between two rotations

In this section, Equation (18) will be proven to be a well-defined distance definition in *SO*(3).

First, the maximum definition of distance between two rotations *R*_**q**1_, *R*_**q**2_ ∈ *SO*(3), i.e., Equation (18), can be attained as *S*^2^ is compact. Thus, the use of the maximum operator is appropriate in Equation (18).

Second, it will be proven below that the three basic requirements of the distance definition in mathematics, i.e., the symmetry property, the positive definitiveness property, and the triangle inequity property, are satisfied for Equation (18).

The symmetry property is such that

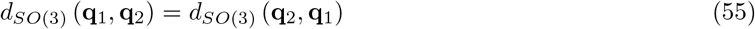

holds for all **q**_1_, **q**_2_ ∈ *SO*(3). It is proven as follows.

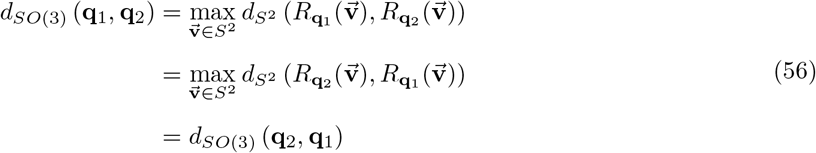

as 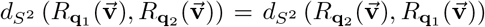 holds, because *d*_*S*^2^_, i.e, distance in *S*^2^, also has symmetry property.

The positive definiteness property contains two parts. The first part is that

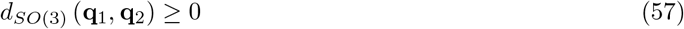

holds for all *R*_**q**_1__, *R*_**q**_2__ ∈ *SO*(3), which can be derived by 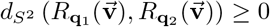 for any 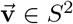, as *d*_*S*^2^_ is the distance in *S*^2^, which also has a positive definiteness property. The second part is that

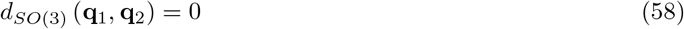

holds when and only when *R*_**q**_1__ = *R*_**q**_2__. This statement is proven as follows. When 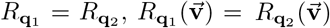 for any 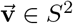, which derives 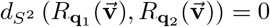 for any 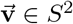, resulting *d*_*SO*(3)_ (*R*_**q**_1__, *R*_**q**_2__) = 0. Inversely, when *d*_*SO*(3)_ (*R*_**q**_1__, *R*_**q**_2__) = 0, for any 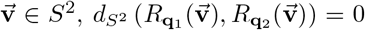, i.e., 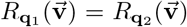; in other words, *R*_**q**_1__ = *R*_**q**_2__.

The triangle inequity property is that

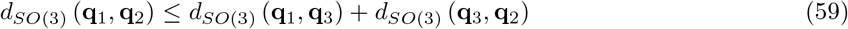

holds for all *R*_**q**_1__, *R*_**q**_2__, *R*_**q**_1__ ∈ *SO*(3). It is proven as follows.

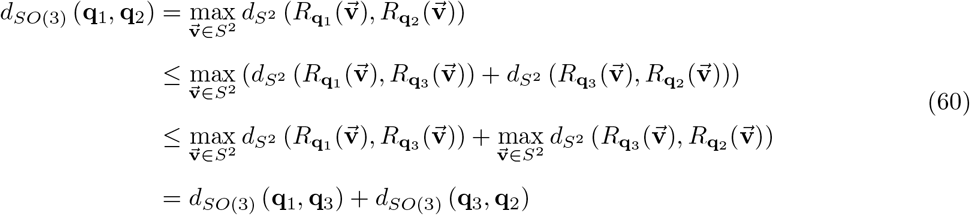

as 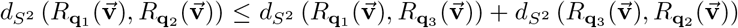 holds because *d*_*S*^2^_, i.e., the distance in *S*^2^ also has a triangle inequity property.

We have proven the suitability of Equation (18) as a distance definition between two rotations.

### Appendix V-II Rotation distance calculated using unit quaternions

In this section, Equation (19) is derived as a method to calculate the distance in *SO*(3) defined by Equation (18) using the unit quaternion. This proof contains two steps. First, it will be shown that the distance between *R*_**q**_1__ and *R*_**q**_2__ in *SO*(3) using Equation (18) equals the angle of rotation of 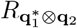. Second, 2 cos^−1^ (|**q**_1_ · **q**_2_|), which is obtained by Equation (19) will also be shown to be equal to the angle of rotation of 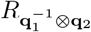.

The first step, i.e., *d*_*SO*(3)_(**q**_1_, **q**_2_) in Equation (18) equals the angle of rotation of 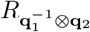, is proven as follows.

*d*_*SO*(3)_ has the following properties. The right multiplication of the same rotation *R*_**q**_ maintains the distance such that

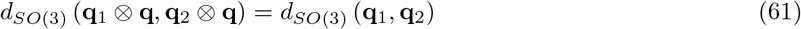

holds for all *R*_**q**_1__, *R*_**q**_2__ and *R*_**q**_. The left multiplication of the same rotation *R*_**q**_ also maintains the distance such that

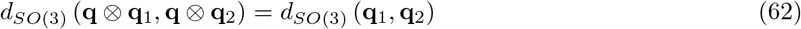

holds for all *R*_**q**_1__, *R*_**q**_2__ and *R*_**q**_.

The right multiplication distance-keeping property, i.e., Equation (61), holds as

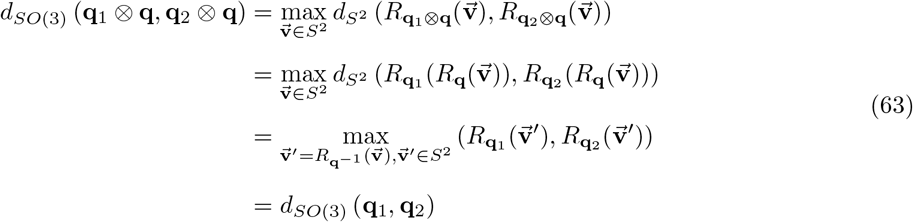

by changing the variable.

The left multiplication distance-keeping property, i.e., Equation (62), holds as

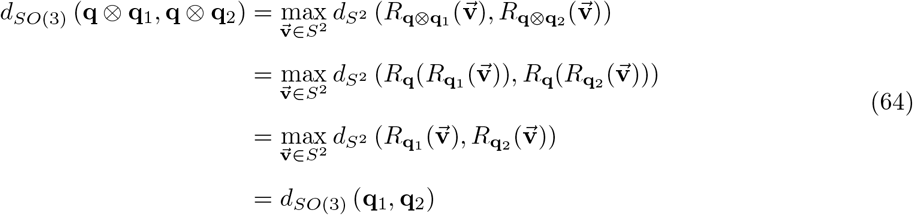

because the distance on the unit sphere *S*^2^ has the following property

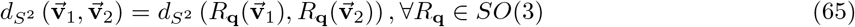

because the spatial rotation preserves the inner product, i.e., Equation (43).

With the properties of the distance on *SO*(3) described in Equation (61) and Equation (62), more properties can be derived as follows.

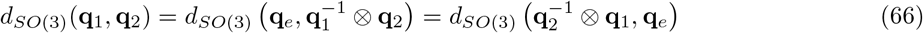

holds for any *R*_**q**_1__, *R*_**q**_2__ ∈ *SO*(3), where **q**_*e*_ represents the still rotation, i.e., the rotation that does not move any vector. Moreover,

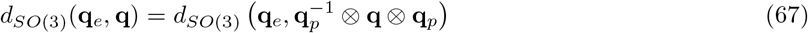

holds for any *R*_**q**_, *R*_**q**_*p*__ ∈ *SO*(3).

For any *R*_**q**_ ∈ *SO*(3), there exists a rotation *R*_**q**_*p*__ ∈ *SO*(3) such that the corresponding rotation matrix of 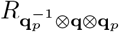 is [3]

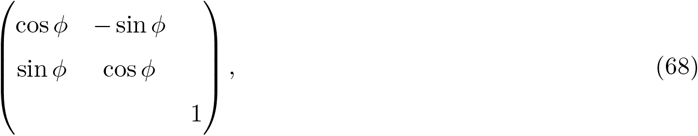

where *ϕ* is the rotation angle of *R*_**q**_. Thus, from Equation (66), Equation (67), and Equation (68), there exist *R*_**q**_*p*__ and *ϕ* ∈ [−*π, π*) such that

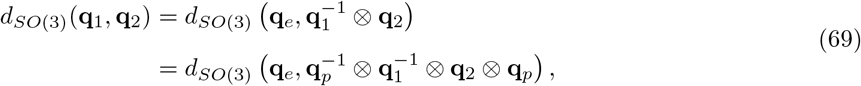

where the corresponding rotation matrix of 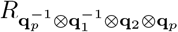 is 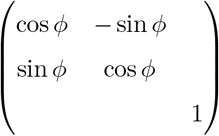, and *ϕ* is the angle of rotation of 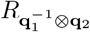.

From the definition of the distance in *S*^2^, i.e., Equation (17), and the definition of the distance in *SO*(3), i.e, Equation (18), Equation (69) is converted to

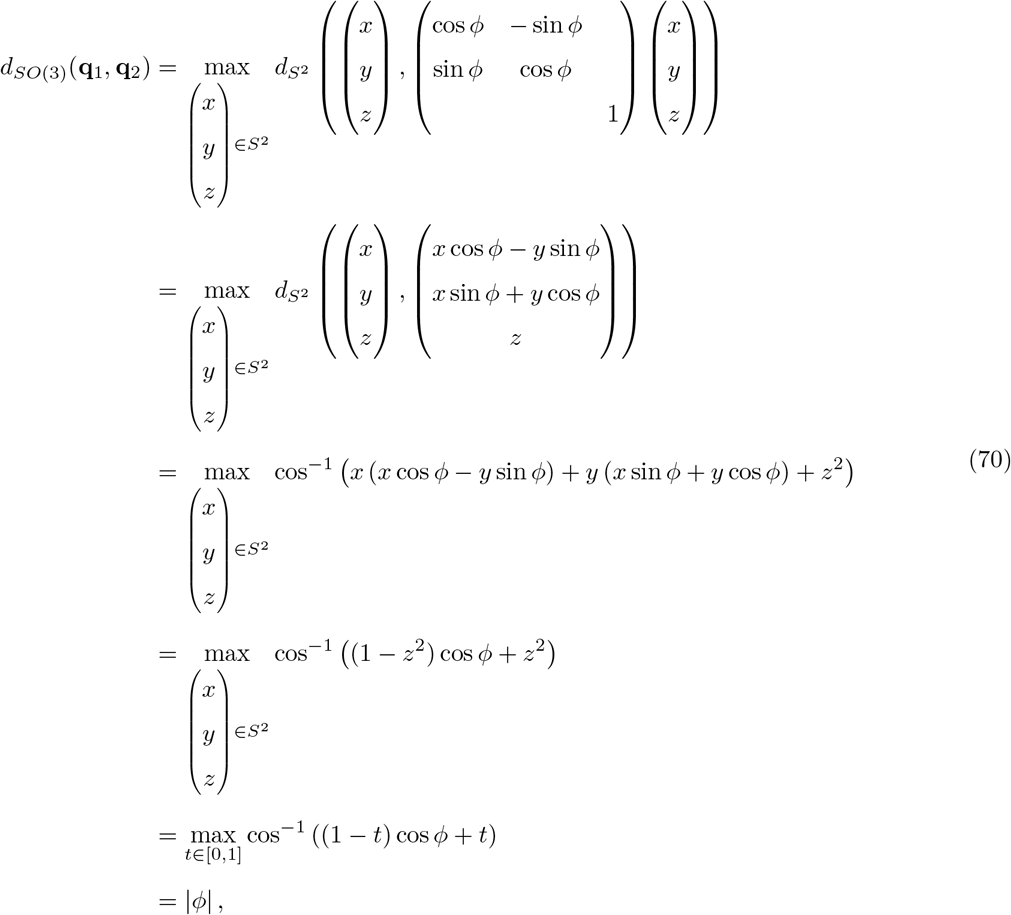

that has a ranges [0, *π*]. By far, the distance *d*_*SO*(3)_ (**q**_1_, **q**_2_) has been shown to be equal to the angle of rotation of 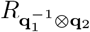.

The second step, i.e., 2cos^−1^ (|**q**_1_ · **q**_2_|) in Equation (19) will also be shown to be equal to the angle of rotation of 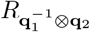.

As the unit quaternion multiplication has this property[4]

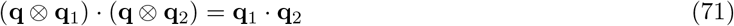

by choosing 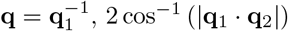 is derived into

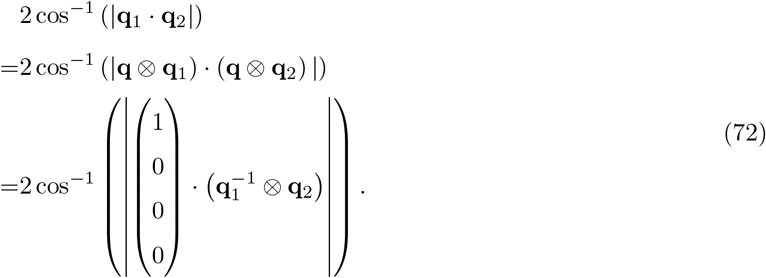

As the rotation angle of 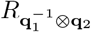 is *ϕ*, fromy Equation (10), 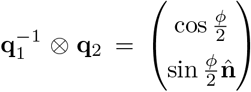 or 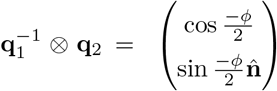, where unit vector 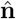 is the rotation axis. Thus, using Equation (72), the second step is proven as

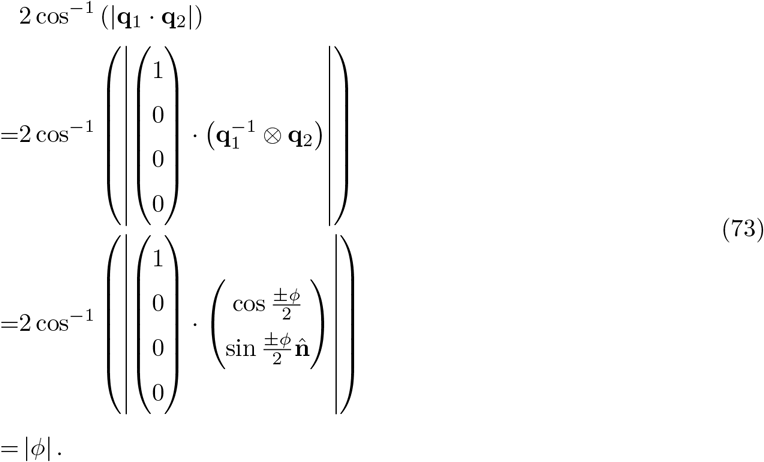

By far, the distance *d*_*SO*(3)_(*R*_**q**_1__, *R*_**q**_2__) defined in Equation (18) and 2cos^−1^ (|**q**_1_ · **q**_2_|) in Equation (19) have both been proven to be the rotation angle of 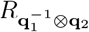. Thus, Equation (19) has been proven.

### Appendix V-III Rotation distance as upper bound of direction change

In this section, we prove that for a 3D object rotated by two rotations *R*_**q**_1__ and *R*_**q**_2__, the distance between the orientations represented by their projection direction, i.e., 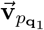 and 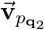, is no larger than the distance *d*_*SO*(3)_ (*R*_**q**_1__, *R*_**q**_2__).

By choosing 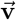 in Equation (18) as 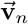 in Equation (12),

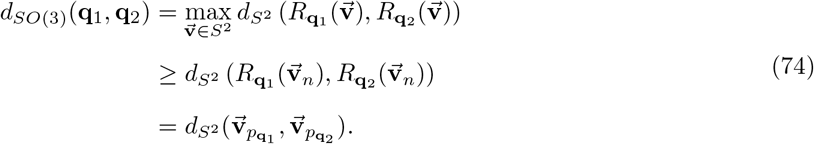

Thus, the distance between *R*_**q**_1__ and *R*_**q**_2__ has been proven to be the upper bound of their projection direction difference.

## Appendix VI Projective arithmetic mean as an approximation of geometric mean

In this section, the projective arithmetic mean will be proven to be an approximation of the geometric mean for a given set of rotations {*R*_**q**_*i*__}_1≤*i*≤*N*_ and their weights {*w_i_*}_1≤*i*≤*N*_.

The projective arithmetic mean is defined as[5]

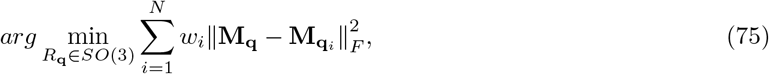

where **M_q_** represents the rotation matrix of rotation *R*_**q**_, and ||·||_*F*_ represents the Frobenius norm[6]. From Equation (19), ||**M_q_** - **M**_**q**_*i*__||_*F*_ is related to the distance defined in Equation (18)[7], such that

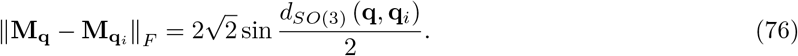

As 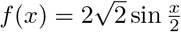 is a monotonic increasing function where *x* ranges [0, *π*), the projective arithmetic mean is an approximation of the geometric mean to some degree.

## Appendix VII Selection of covariance matrix that causes the ACG distribution to reach a maximum probability density at a given unit quaternion

In this section, we introduce a convenient method for selecting the covariance matrix **A** in the ACG distribution to make the *ACG*(**A**) attain a maximum probability density at a given unit quaternion **q**. As stated in Section 4.3.1, **A** should be a positive-definite symmetric matrix with **q** as the principal eigenvector.

Assuming 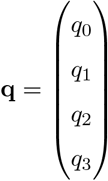, we can choose **A** as

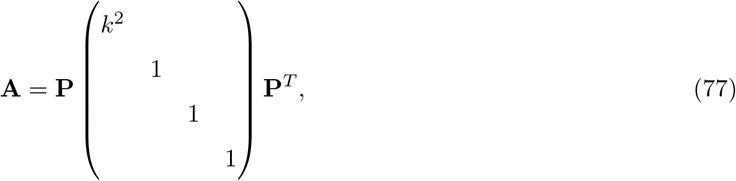

where

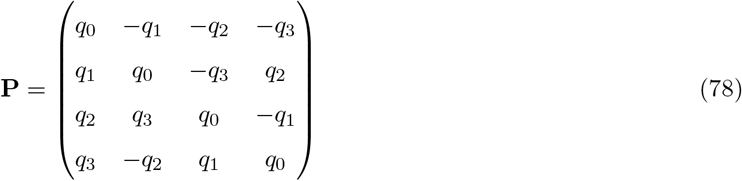

is an orthogonal matrix and *k* > 1.

As, 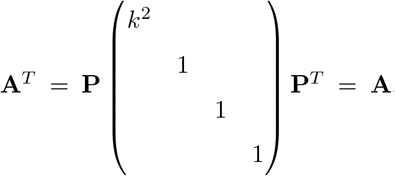, **A** is symmetric. Moreover, as *k* > 1, from the principle of eigenvalue decomposition[3], **q** is the principal eigenvector of **A**.

## Appendix VIII Probability density of unit vector generated by rotations with ACG distribution

In this section, we discuss the distribution of a unit vector rotated by rotations sampled from an ACG distribution with a special covariance matrix 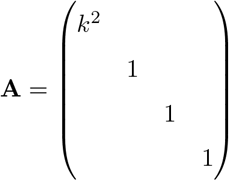, where *k* > 1. For a vector **v**_0_ and a unit quaternion **q** sampled from *ACG*(**A**), the rotated vector 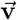 is calculated as 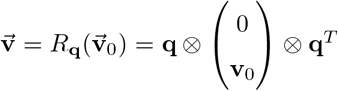. We denote the probability density of this distribution of 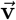 as 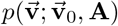.

The concept of the deviation of 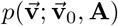 is to use the principle of marginal probability density, such that

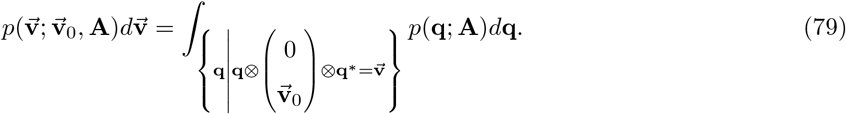

We will prove that 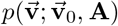 can be calculated as

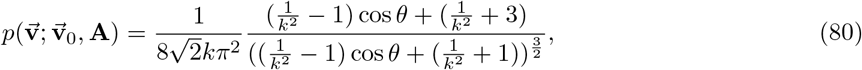

where 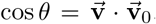, i.e., *θ* is the angular distance between 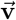 and 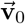. From Equation (80), we can also conclude that 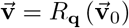 obeys a distribution invariant under rotation around 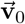. Moreover, by converting Equation (80) to

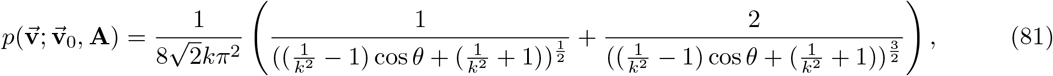

we can conclude that 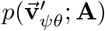 monotonically decreases when *θ* increases as *k* > 1, which means that 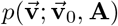, i.e., the probability density of 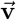 reaches a maximum at 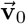.

In the first part of this proof, we will specify 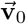 to be (0, 0,1)^*T*^ and prove the above statement. Then, in the second part of this proof, we will extend 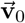 from (0,0,1)^*T*^ to an arbitrary unit vector, while the above statement will still hold.

For the first part, 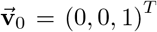. We use Euler angles to parameterize spatial rotations. Considering *ZY Z* proper Euler angles *ϕ, θ*, and *ψ*, where *ϕ* ∈ [0, 2*π*), *θ* ∈ [0, *π*], and *ψ* ∈ [0, 2*π*), rotation *R*_*ϕ,θ,ψ*_ is given by

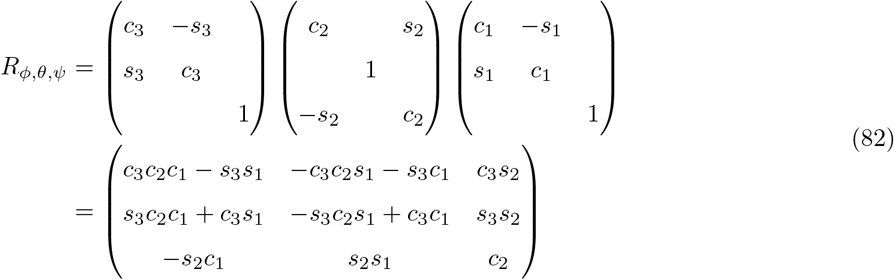

where *c*_1_ = cos *ϕ*, *s*_1_ = sin *ϕ*, *c*_2_ = cos*θ*, *s*_2_ = sin*θ, c*_3_ = cos *ψ*, and *s*_3_ = sin*ψ*. Let *R*_ϕ,θ,ψ_ act on 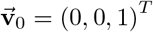,

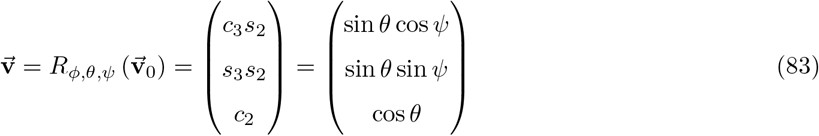

we can observe that 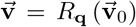 is independent of *ϕ*. In other words, 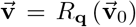 obeys a distribution invariant under rotation around 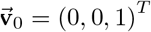.

Recall that the rotation of *ZY Z* Euler angles corresponds to a unit quaternion given by Equation (51), and the Haar measure under these coordinates is 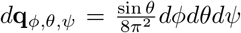[8]. Therefore, based on the ACG probability density function described in Equation (31), the probability density of the ACG distribution using Euler angles is

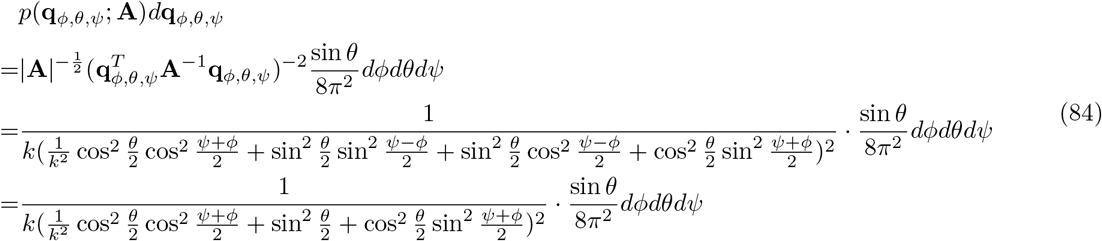

where *p*(**q**_*ϕ,θ,ψ*_; **A**) is the probability density of the ACG distribution with covariance matrix as **A**.

By substituting Equation (84) into (79),

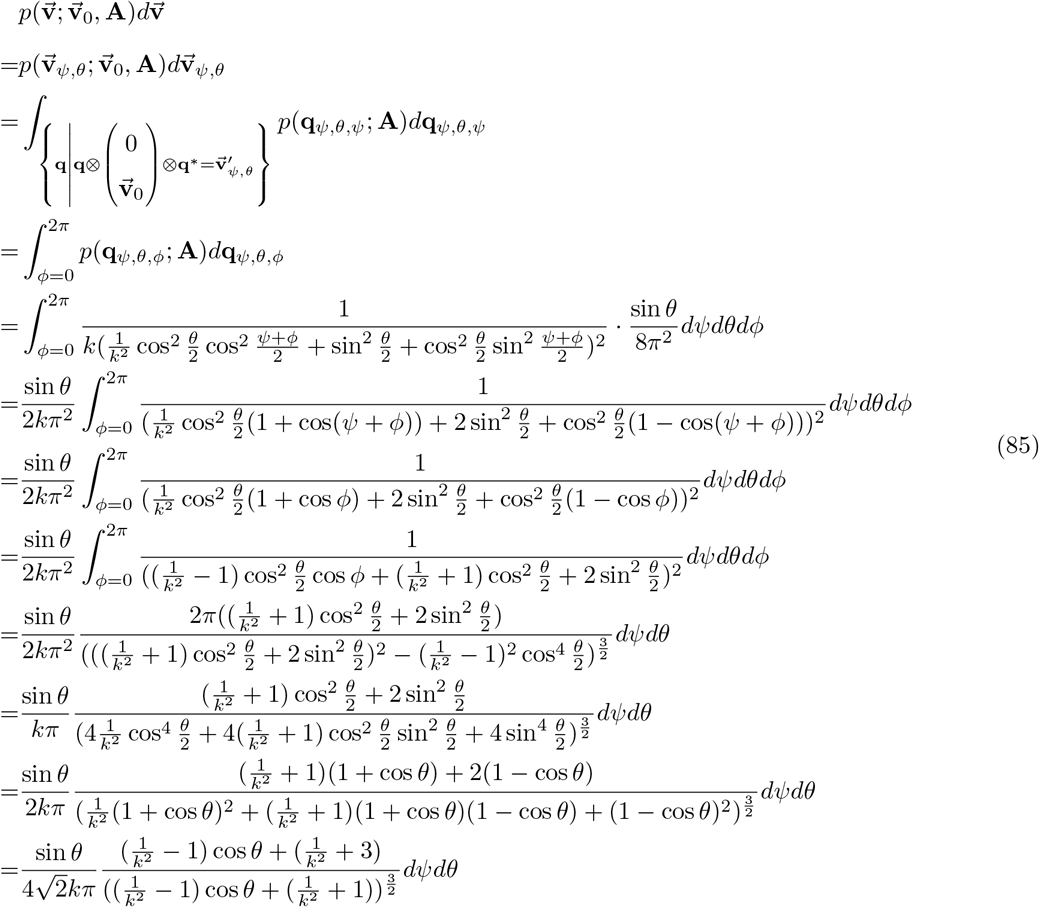

In *S*^2^, 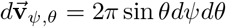. Thus,

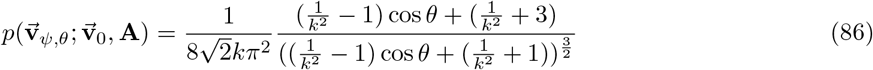

Therefore, we have proven Equation (80) when 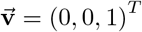.

In the second part, we will extend 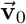 from (0, 0,1)^*T*^ to an arbitrary unit vector, while the above statement will hold. Assume that 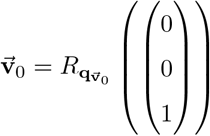, that is, 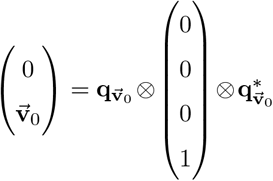, Equation (79) can be further derived as

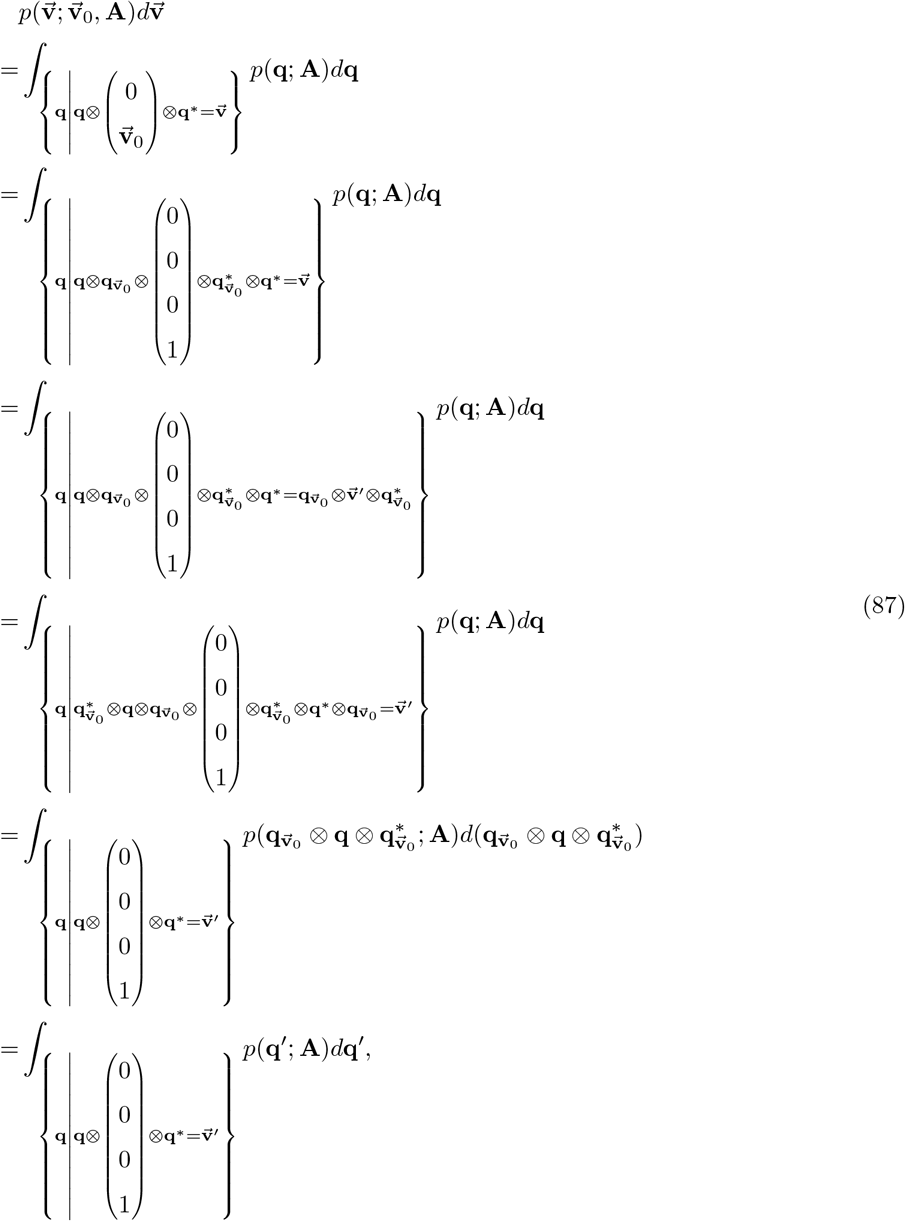

where 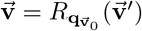 and 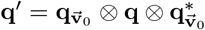.

Note that *d*_**q**_ is the Haar measure on *S*^3^, so[9]

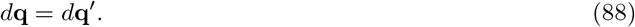

Moreover,

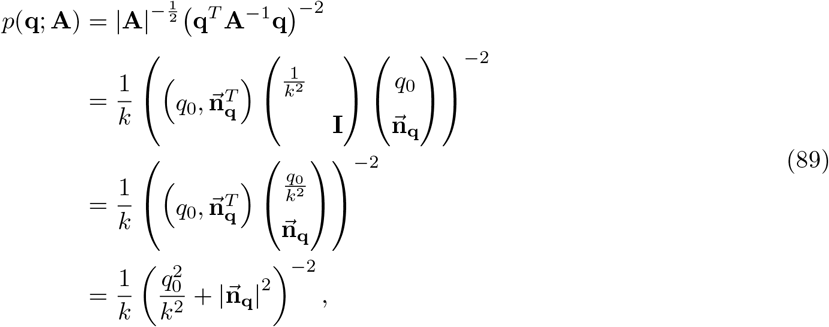

where 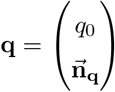. As

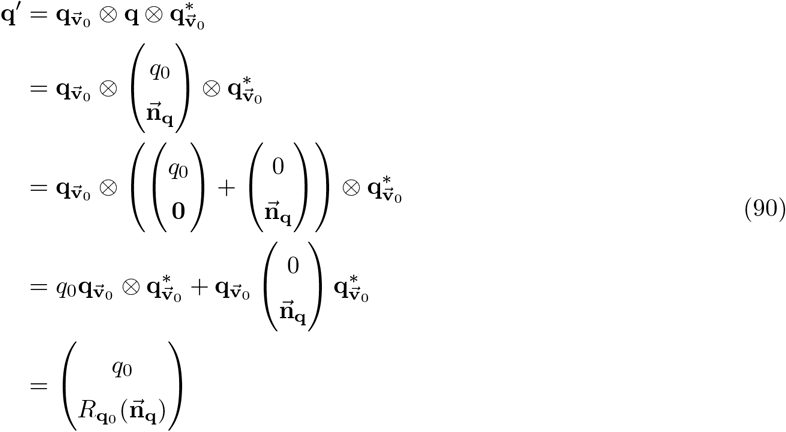

and 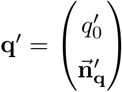, we can conclude that 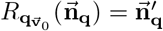. As 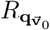 is a spatial rotation, 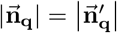. Thus, we can conclude from Equation (89) that

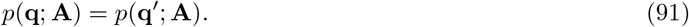

By substituting Equation (88) and Equation (91) into Equation (87), we can conclude that

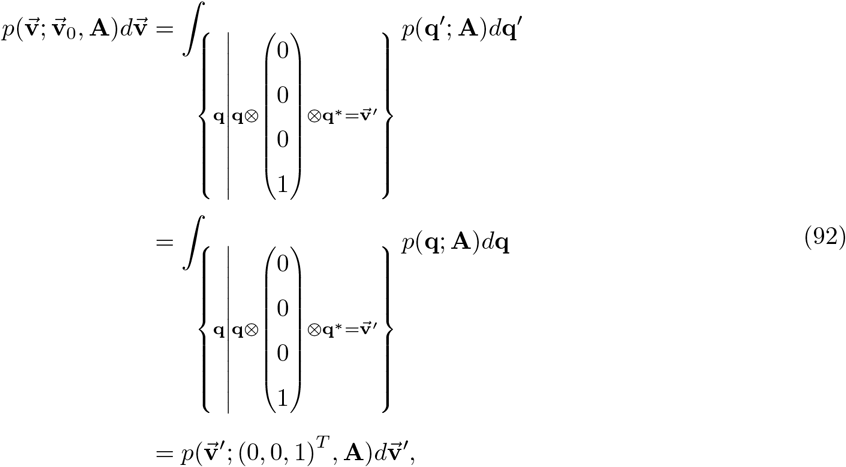

where 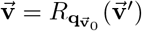. As

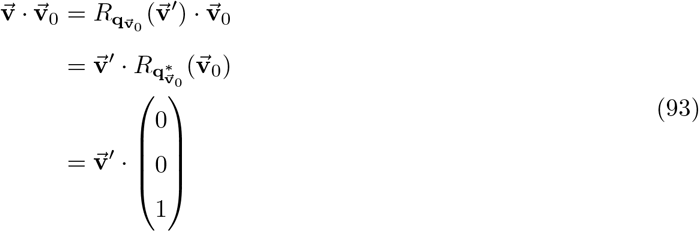

and 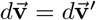, Equation (86) can be converted into

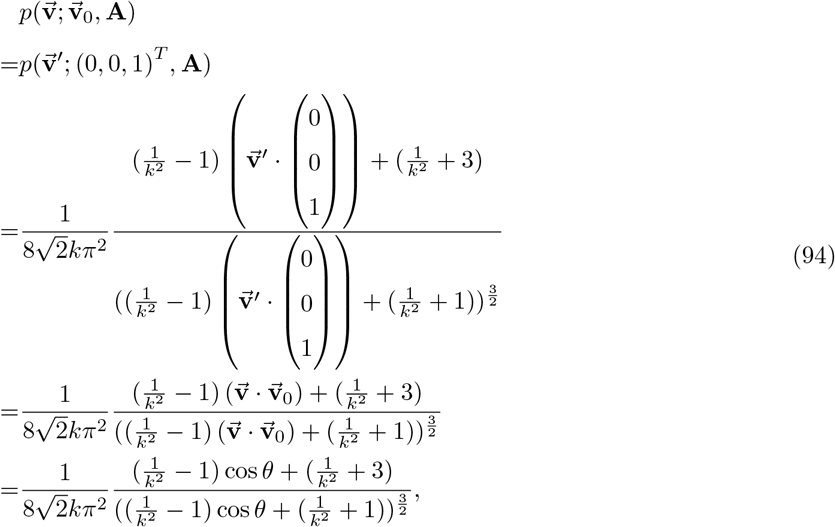

where 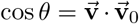. Up to the present, we have proven that Equation (80) still holds when 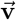 is an arbitrary unit vector.

## Appendix IX Confidence area of a unit vector rotated by rotations from ACG distribution with a special covariance matrix

In this section, we still follow the assumptions and notations in Appendix VIII. We will prove that the method employed to calculate the confidence area using Equation (37) holds.

As 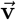 is in *S*^2^, which means that 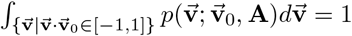, Equation (36) is equivalent to

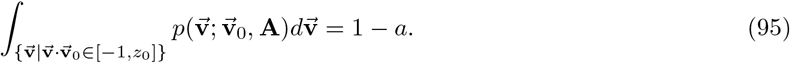

By substituting Equation (35) into Equation (95),

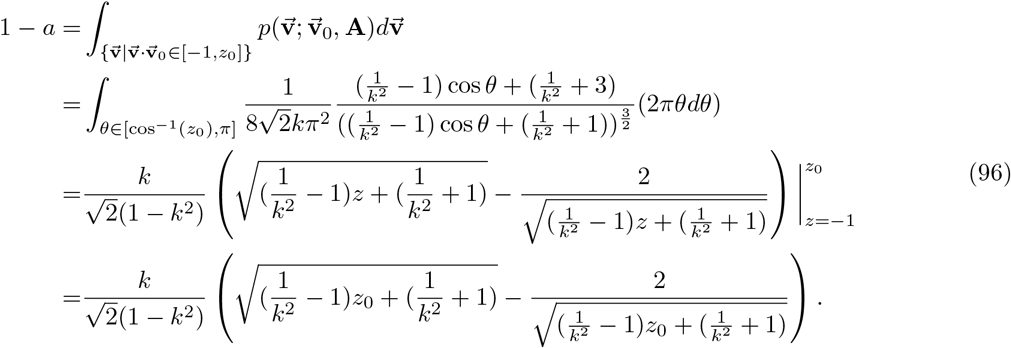

For simplicity, we denote 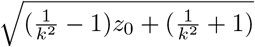 as 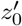 and 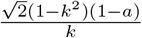 and *a*′; thus,

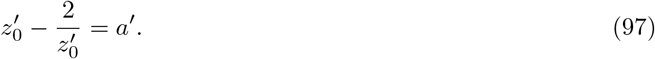

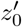 is the positive solution of 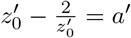, as in

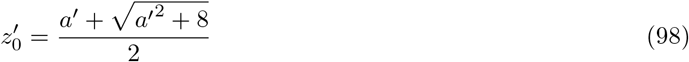

from which *z*_0_ can be determined as

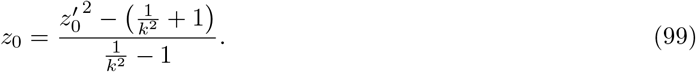

We have thus proven that the method employed to calculate the confidence area using Equation (37) holds.

## Appendix X Approximation formula used to calculate confidence area of a unit vector rotated by rotations from ACG distribution using a special covariance matrix

In this section, we still follow the assumptions and notations in Appendix VIII and Appendix IX. We prove that the reciprocal of angular precision, i.e., 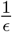, is asymptotically linear to *k*, and that the asymptote is

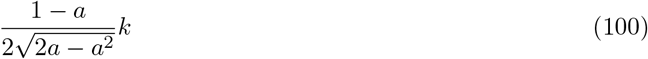

to which 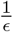 trends when *k* →∞.

The proof is as follows. From Equation (37), it can be seen that

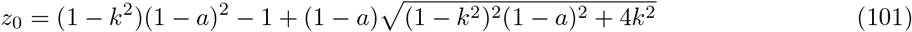

Note that when *k* →∞,

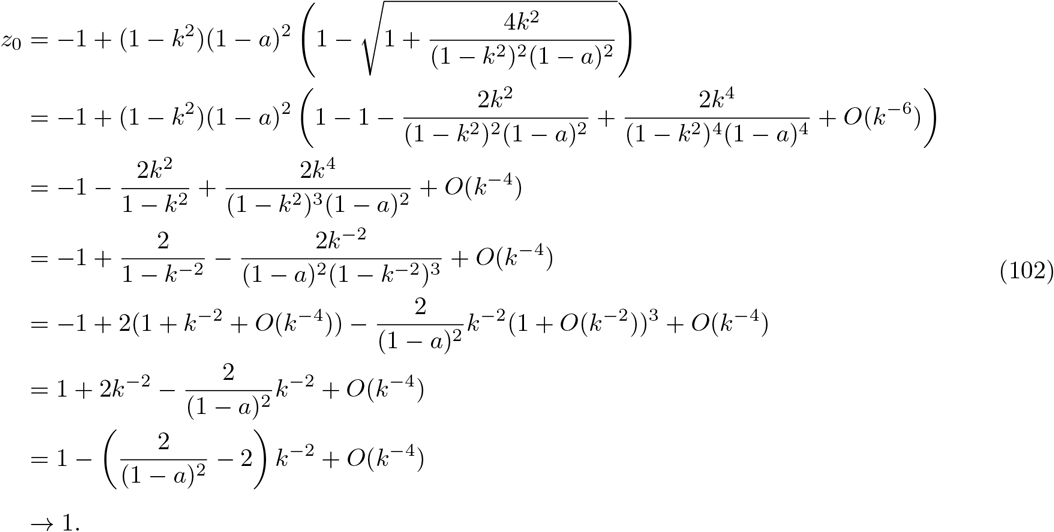

Hence, *θ*_0_ → 0 and 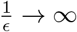. We aim to show that when *k* → ∞, 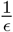 grows linearly in *k*. We have

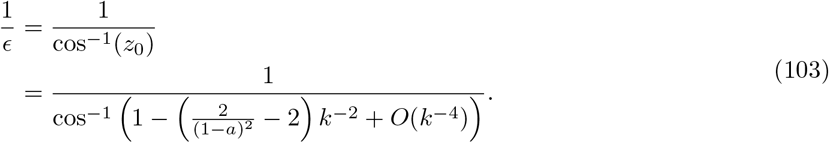

By Puiseux expansion, cos^−1^(1 – *x*) is

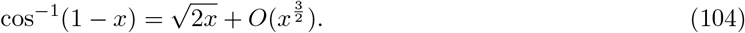

It follows that

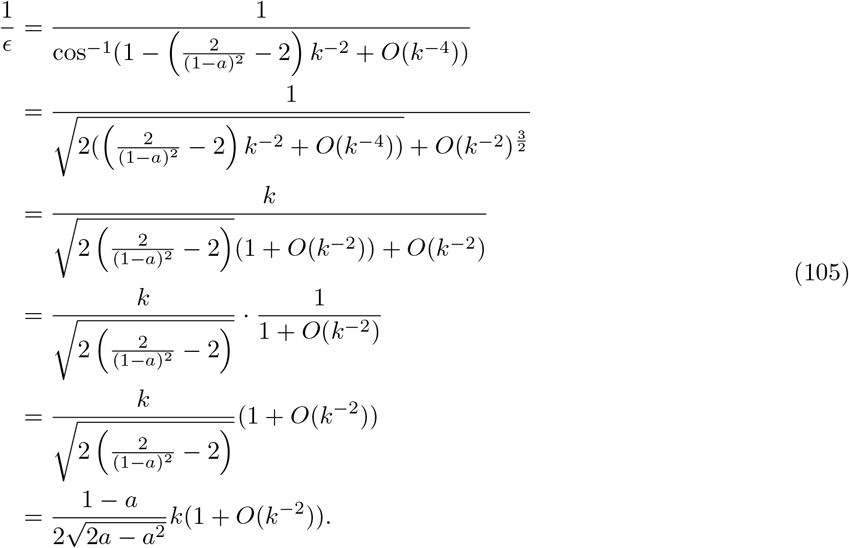

Hence, we have proven that the reciprocal of angular precision, i.e., 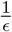, is asymptotically linear to *k*, and the asymptote is 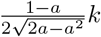, to which 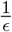 trends when *k* → ∞.

## Appendix XI Special form of ACG distribution inference

In this section, we will prove that the iterative method for determining the maximum-likelihood approximation of *k* in the special-form ACG distribution covariance matrix **A** holds.

We denote *k*^2^ as *ξ*, and 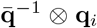 as 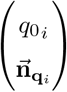. We imitate the method of this reference[10] to find the maximum-likelihood estimate of parameter *k* for *ACG*(**A**), where 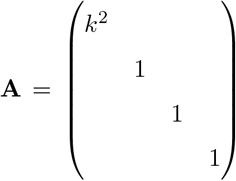. The likelihood function for *ξ* is

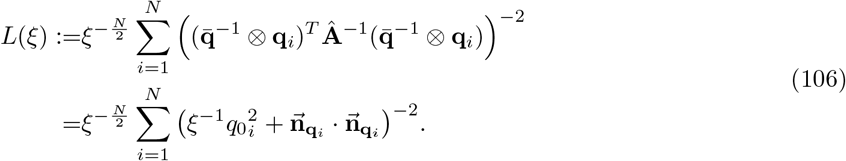

The maximum-likelihood estimate *ξ* must satisfy

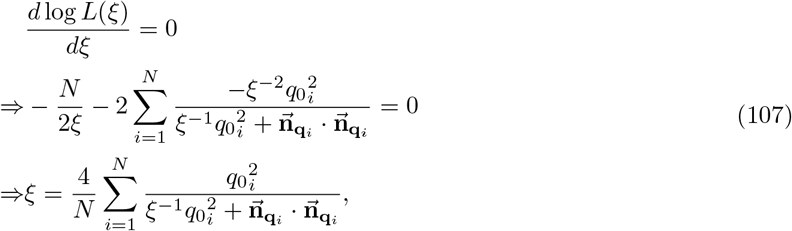

where *ξ* = *k*^2^. Thus, we have proven Equation (39) as an iteration formula for solving *k*.

